# Carryover effects of temperature and pCO_2_ across multiple Olympia oyster populations

**DOI:** 10.1101/616375

**Authors:** Laura H Spencer, Yaamini R Venkataraman, Ryan Crim, Stuart Ryan, Micah J Horwith, Steven B Roberts

**Affiliations:** University of Washington, School of Aquatic and Fishery Sciences, 1122 NE Boat St, Seattle, WA 98105, United States; Puget Sound Restoration Fund, 8001 NE Day Rd W, Bainbridge Island, WA 98110, United States; Washington State Department of Natural Resources, 1111 Washington St SE, MS 47027, Olympia, WA 98504, United States

**Keywords:** Ostrea lurida, acidification, pH, warming, winter, reproduction, phenology, intergenerational, transgenerational, climate change

## Abstract

Predicting how populations will respond to ocean change across generations is critical to effective conservation of marine species. One emerging factor is the influence of parental exposures on offspring phenotype, known as intergenerational carryover effects. Parental exposure may deliver beneficial or detrimental characteristics to offspring that can influence larval recruitment patterns, thus shaping how populations and community structure respond to ocean change. Impacts of adult exposure to elevated winter temperature and pCO_2_ on reproduction and offspring viability were examined in the Olympia oyster (*Ostrea lurida*) using three populations of adult, hatchery-reared *O. lurida,* plus an additional cohort spawned from one of the populations. Oysters were sequentially exposed to elevated temperature (+4°C, at 10°C), followed by elevated pCO_2_ (+2204 µatm, at 3045 µatm) during winter months. Male gametes were more developed after elevated temperature exposure and less developed after high pCO_2_ exposure, but there was no impact on female gametes or sex ratios. Oysters previously exposed to elevated winter temperature released larvae earlier, regardless of pCO_2_ exposure. Those exposed to elevated winter temperature as a sole treatment released more larvae on a daily basis, but when also exposed to high pCO_2_ there was no effect. These combined results indicate that elevated winter temperature accelerates *O. lurida* spermatogenesis, resulting in earlier larval release and increased production, with elevated pCO_2_ exposure negating effects of elevated temperature. Altered recruitment patterns may therefore follow warmer winters due to precocious spawning, but these effects may be masked by coincidental high pCO_2_. Offspring were reared in common conditions for one year, then deployed for three months in four estuarine bays with distinct environmental conditions. Offspring of parents exposed to elevated pCO_2_ had higher survival rates in two of the four bays. This carryover effect demonstrates that parental conditions can have substantial ecologically relevant impacts that should be considered when predicting impacts of environmental change. Furthermore, Olympia oysters may be more resilient in certain environments when progenitors are pre-conditioned in stressful conditions. Combined with other recent studies, our work suggests that the Olympia may be more equipped than other oysters for the challenge of a changing ocean.

## Introduction

The repercussions of ocean warming and acidification on marine invertebrate physiology are complex, but significant recent advances indicate that larval stages of marine taxa are particularly vulnerable (Byrne & Przeslawski, 2013; Kurihara, 2008; Przeslawski, Byrne, & Mellin, 2015). Understanding how shifting conditions will influence larval recruitment patterns is critical to predicting changing population dynamics, and thus community structure. One emerging consideration is whether larval stages benefit from ancestral exposures, based on evidence that memory of environmental stressors can be transferred between generations through non-genetic inheritance (reviewed in Perez & Lehner, 2019; Donelson *et al*. 2018; Eirin-Lopez & Putnam, 2019; Ross, Parker, & Byrne, 2016). Beneficial, or positive, carryover effects may be important acclimatory mechanisms for marine organisms facing rapid change, particularly those that evolved in dynamic environments like estuaries and the intertidal (Donelson, Salinas, Munday, & Shama, 2018; Gavery & Roberts, 2014). These carryover effects are defined as transgenerational when they persist in generations that were never directly exposed. Intergenerational, or parental, effects may be due to direct exposure as germ cells (Perez & Lehner, 2019). Trans- and intergenerational carryover effects are increasingly reported across marine phyla, including Cnidaria (*e.g.* Putnam & Gates, 2015), Echinodermata (*e.g.* Clark *et al*., 2019), Mollusca (*e.g.* Parker *et al*. 2015), Arthropoda (*e.g.* Thor & Dupont, 2015), and Chordata (Review: Munday 2014).

A foundational series of studies on the Sydney rock oyster (*Saccostrea glomerata*) provide strong evidence for intergenerational carryover effects in bivalves, an ecologically and economically important group of taxa (Dumbauld, Ruesink & Rumrill, 2009). Adult *S. glomerata* exposed to high pCO_2_ produced larger larvae that were less sensitive to high pCO_2_, and the effect persisted in the successive generation (Parker *et al*., 2012, 2015). In the presence of secondary stressors, however, parental high pCO_2_ exposure rendered larvae more sensitive (Parker *et al*., 2017). Intergenerational carryover effects are increasingly documented in larvae across other bivalve species, and are beneficial in the mussels *Mytilus chilensis* (Diaz *et al*., 2018) and *Mytilus edulis* (but not juveniles) (Kong *et al*., 2019; Thomsen *et al*., 2017), and detrimental in the clam *Mercenaria mercenaria*, the scallop *Argopecten irradians* (Griffith & Gobler, 2017), and the oyster *Crassostrea gigas* (Venkataraman, Spencer, & Roberts, 2019).

These preliminary studies provide strong evidence for intergenerational carryover effects in bivalves, but the body of work is still narrow in scope. Nearly all studies have exposed parents to stressors during denovo gamete formation (gametogenesis). For many temperate bivalve species, this occurs seasonally in the spring (Bayne, 1976). Yet, challenging periods of acidification and warming can occur during other times of the year (Evans, Hales, & Strutton, 2013; Joesoef, Huang, Gao, & Cai, 2015; McGrath, McGovern, Gregory, & Cave, 2019). The most corrosive carbonate environment in the Puget Sound estuary in Washington State, for example, commonly occurs in the winter when many species are reproductively inactive, while favorable conditions are in the spring when gametogenesis coincides with phytoplankton blooms (Pelletier, Roberts, Keyzers, & Alin, 2018). Thus, adult exposure to severely corrosive conditions during gametogenesis may not represent the natural estuarine system. To our knowledge, only one study has assessed carryover effects of exposure to acidification before reproductive conditioning in a bivalve, the oyster *C. gigas*, and found negative maternal carryover effects on larval survival (Venkataraman, Spencer, & Roberts, 2019), indicating that pre-gametogenic exposure also matters. No studies have yet attempted to examine intergenerational carryover effects of combined winter warming and acidification in bivalves. To best predict whether intergenerational carryover effects will be beneficial or detrimental, it is also crucial to understand how warming and acidification will impact fertility and reproductive phenology. Temperature is a major driver of bivalve reproduction, and modulates gametogenesis (Joyce, Holthuis, Charrier, & Lindegarth, 2013; Maneiro, Pérez-Parallé, Pazos, Silva, & Sánchez, 2016; Oates, 2013), influences sex determination (Santerre *et al*., 2013) and, in many species, triggers spawning (Fabioux, Huvet, Le Souchu, Le Pennec, & Pouvreau, 2005) (alongside other factors such as photoperiod, nutrition, lunar/tidal phases). Year-round warming may result in unexpected impacts to larval competency resulting from changes to reproduction. For instance, some temperate bivalve species have a thermal threshold for gametogenesis and enter a period of reproductive inactivity, or “quiescence”, which is believed necessary for successive spawning (Giese, 1959; Hopkins, 1937; Loosanoff, 1942).

Warmer winters brought on by global climate change (IPCC, 2013, 2019) may therefore shift species’ reproductive cycles to begin earlier, or eliminate seasonality altogether, resulting in poorly provisioned or ill-timed larvae (Chevillot *et al*., 2017). Such impacts were clearly demonstrated using a long-term dataset (1973-2001) of estuarine clam *Macoma balthica* reproduction and temperature. Mild winters and earlier springs resulted in low fecundity, earlier spawning, and poor recruitment, which was largely explained by a phenological mismatch between spawning and peak phytoplankton blooms (Philippart *et al*., 2003). The impacts of winter acidification on estuarine bivalve reproduction are less predictable. The few studies to date show that high pCO_2_ delays gametogenesis in the oysters *Crassostrea virginica* and *S. glomerata* (Boulais *et al*., 2017; Parker *et al*., 2018), but both studies exposed oysters during gametogenesis. Acidification during the winter months could increase energetic requirements (Sokolova, Frederich, Bagwe, Lannig, & Sukhotin, 2012), and deplete glycogen reserves that are later utilized for gametogenesis in the spring (Mathieu & Lubet, 1993), but this hypothesis has yet to be tested.

The purpose of this study was to assess whether warmer, less alkaline winters will affect fecundity and offspring viability in the Olympia oyster, *Ostrea lurida*. The Olympia is native to the Pacific coast of North America (McGraw, 2009). Overharvest and pollution devastated populations in the early 1900s, and today 2-5% of historic beds remain (Blake & Bradbury, 2012; Polson & Zacherl, 2009). Restoration efforts are afoot, but *O. lurida* populations continue to struggle, and may be further challenged by changing conditions (Barton, Hales, Waldbusser, Langdon, & Feely, 2012; Feely, Klinger, Newton, & Chadsey, 2012; Feely, Sabine, Hernandez-Ayon, Ianson, & Hales, 2008). For instance, large interannual variability in larval recruitment and frequent recruitment failures were recently reported (Wasson *et al*., 2016; Kimbro, White & Grosholz, 2019). This variability is presumably related to inconsistent spawning success, larval survival, and retention, and governed predominantly by local conditions (Kimbro, White & Grosholz, 2019). It is unknown how the intensity, timing, and duration of local environmental conditions can predict recruitment failure (Wasson *et al*., 2016). If winter conditions significantly influence recruitment through direct changes to adult reproductive capacity or timing, or indirect changes through parental carryover effects, population densities and distributions will inevitably shift with conditions.

Another consideration in this study was the genetic composition of test organisms. *Ostrea lurida* exhibits varying phenotypes among distinct populations (Silliman, 2019), which can influence their sensitivity to environmental stressors (Bible & Sanford, 2016; Bible, Evans & Sanford, 2019). Indeed, the two groups to measure the response of *O. lurida* larvae to ocean acidification found contrasting results ⎼ no effect (Waldbusser *et al*., 2016), and slower growth (Hettinger *et al*., 2012, 2013) ⎼ possibly a result of local adaptation. The source population used for experimental studies may therefore be a critical factor influencing climate-related findings.

Furthermore, testing genetically diverse organisms could reveal cryptic genetic variation, alleles that confer stress resilience only under certain settings (Paaby & Rockman, 2014; Bitter *et al*., *preprint*), which has implications for how wild populations are restored. Therefore, we tested three phenotypically distinct Puget Sound populations (Heare, Blake, Davis, Vadopalas, & Roberts, 2017; Silliman, Bowyer, & Roberts, 2018), which were hatchery-reared in common conditions to adulthood, to account for intraspecific variation while controlling for within-generation carryover effects (Hettinger *et al*., 2012, 2013).

Our study is the first to assess the combined effects of elevated winter temperature and pCO_2_ on reproduction, and to explore intergenerational carryover in an *Ostrea* spp. We exposed adult *O. lurida* to elevated temperature (+4°C), followed by elevated pCO_2_ (+2204 µatm, -0.51 pH). Gonad development, reproductive timing, and fecundity were assessed for the adults in the laboratory, and offspring performance was assessed in the field. Elevated winter temperature was expected to impede gametogenic quiescence, presumably a critical annual event, subsequently reducing larval production. This prediction was in part based on observations of low larval yields in an *O. lurida* restoration hatchery (Ryan Crim, *unpublished*) following the winter 2016 marine heat wave in the Northeast Pacific Ocean (Gentemann, Fewings, & García-Reyes, 2017).

Similarly, we predicted that high pCO_2_ exposure would result in negative impacts due to increased energy requirements for calcification and cellular maintenance. Finally, we predicted that negative impacts would be amplified upon exposure to both conditions. By assessing the effects of winter warming and acidification on reproduction and offspring viability in multiple Olympia oyster populations, we provide an ecologically relevant picture of how the species will respond to ocean change.

## Methods

### Adult oyster temperature and pCO_2_ exposures

Four cohorts of adult *Ostrea lurida* were used in this study. Three of the cohorts were first-generation hatchery-produced (F1) oysters (32.1 ± 5.0 mm), all hatched in Puget Sound (Port Gamble Bay) in 2013 (Heare *et al*., 2017). The broodstock used to produce these F1 oysters were wild, harvested from Fidalgo Bay in North Puget Sound (F), Dabob Bay in Hood Canal (D), and Oyster Bay in South Puget Sound (O-1) (O in Figure 1). These populations are considered phenotypically distinct subpopulations (Heare *et al*., 2017; White, Vadopalas, Silliman, & Roberts, 2017). The fourth cohort (O-2, 21.9 ± 3.3 mm) was second-generation, hatchery-produced in 2015 from the aforementioned Oyster Bay F1 cohort, from a single larval release pulse and thus likely one family (Silliman, Bowyer, & Roberts, 2018). The O-2 cohort was included to examine whether reproductive and offspring traits were consistent across generations of a population, with the O-2 cohort being closely related to each other (siblings) and 2 years younger than the other cohorts. Prior to the experiment, all oysters were maintained in pearl nets in Clam Bay (C) for a minimum of 500 days.

**Figure 1:**
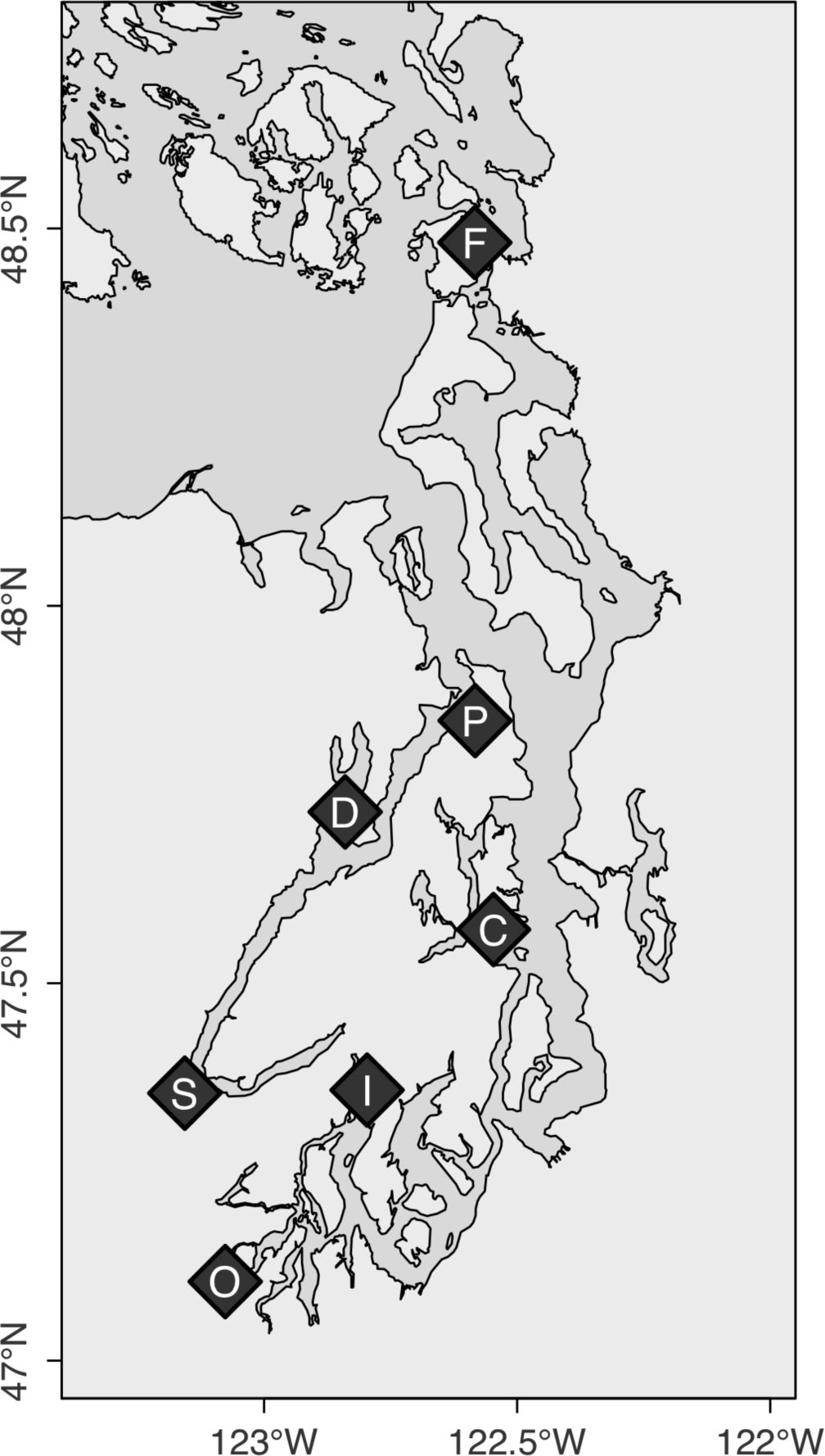
Locations where *O. lurida* populations’ progenitors were collected (F, D, O), where oysters were housed prior to and during the experiment (C), and where offspring were deployed (F, P, S, I): Fidalgo Bay (F), Port Gamble Bay (P), Dabob Bay (D), Clam Bay (C), Skokomish River Delta (S), Case Inlet (I), Oyster Bay (O).

### Temperature treatment

Oysters were moved from Clam Bay (C) to the Kenneth K. Chew Center for Shellfish Research and Restoration for the temperature and pCO_2_ experiments. Oysters were held in one of two temperature regimes (6.1±0.2°C and 10.2±0.5°C) for 60 days beginning December 6, 2016 (Figure 2). The temperatures correspond to historic local winter temperature (6°C) in Clam Bay, and anomalously warm winter temperature (10°C) as experienced during 2014-2016 (Gentemann *et al*., 2017). For the temperature exposure, oysters from each cohort (100 for O-1 and F cohorts, 60 for D, and 300 for O-2) were divided into four bags, two bags per temperature, in two flow-through experimental tanks (50L - 1.2-L/min). Temperature in the 6°C treatment was maintained using an aquarium chiller, and unchilled water was used for the 10°C treatment. Temperatures were recorded continuously with water temperature data loggers.

**Figure 2:**
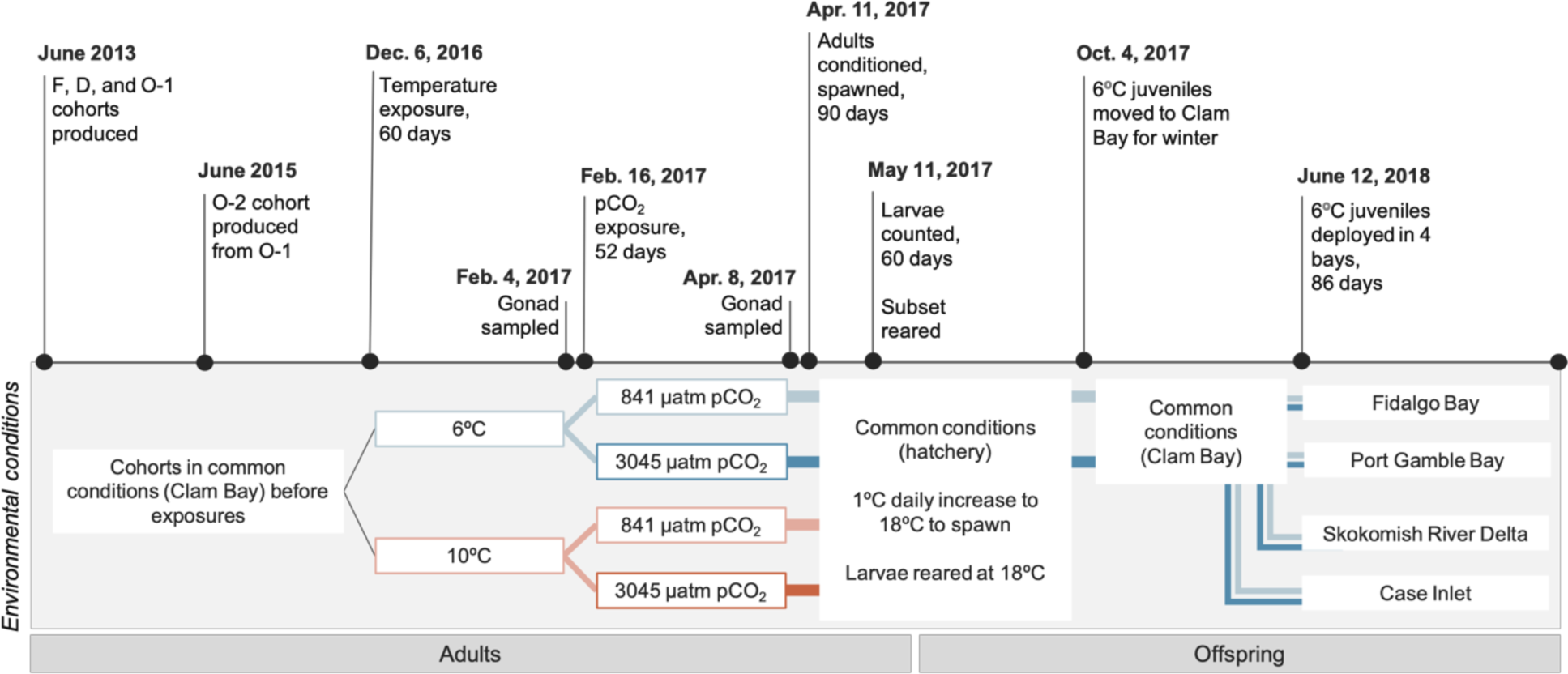
Experimental timeline. Four cohorts of adult *O. lurida* (F, D, O-1, O-2) were sequentially exposed to two winter temperatures (6.1±0.2°C, 10.2±0.5°C) then two pCO_2_ levels (841±85 µatm, 3045±488 µatm). They were returned to ambient pCO_2_ conditions to volitionally spawn. Larvae were collected and reared by cohort x temperature x pCO_2_. Juveniles (∼1 year) from 6°C-Ambient pCO_2_ and 6°C-Low pCO_2_ adults were deployed in 4 bays in Puget Sound.

### High pCO_2_ treatment

A differential pCO_2_ exposure was carried out after the temperature treatment ended. Following a 10-day gradual temperature increase for the 6°C treatment to 10°C, oysters were further divided and held at ambient pCO_2_ (841±85 µatm, pH 7.82±0.02) or high pCO_2_ (3045±488 µatm, pH 7.31 ± 0.02) for 52 days (February 16 to April 8, 2017, Figure 2). Animals were housed in six flow-through tanks (50-L - 1.2-L/min), with three replicate tanks per pCO_2_ treatment and oyster cohort. High pCO_2_ treated water was prepared using CO_2_ injection. Filtered seawater (1µm) first recirculated through a reservoir (1,610-L) with a degassing column to equilibrate with the atmosphere, then flowed into treatment reservoirs (757-L) recirculating through venturi injectors. Durafet pH sensors and a Dual Input Analytical Analyzer monitored pH in treatment reservoirs with readings every 180 seconds. Using solenoid valves, CO_2_ gas was injected through lines at 15 psi in 0.4 second pulses if pH exceeded the 7.22 set point. Water pH was continuously monitored in experimental tanks using Durafet pH sensors, and temperature (10.4 ± 0.4°C) was measured using water temperature data loggers. Twice weekly, water samples (1-L) were collected from experimental tanks, and temperature (°C), salinity (PSU), and pH (mV, converted to pH_T_) were measured immediately using a digital thermometer, conductivity meter, and pH electrode, respectively. Simultaneously, discrete water samples (120-mL) were collected in duplicate from experimental tanks and preserved with HgCl (50-µL) for later total alkalinity measurements using a titrator. Standard pH curves were generated on each sampling day prior to pH measurements using TRIS buffer prepared in-house at five temperatures (Appendix S1: Section S1). Using the `seacarb` library in R, pCO_2_, dissolved organic carbon (DIC), calcite saturation (Ω_calcite_), and aragonite saturation (Ω_aragonite_) were calculated for days 5, 33, and 48 (Appendix S1: Table S1).

**Table 1:** Environmental data during offspring field trial. Environmental data was collected from locations where offspring were deployed for 3 months from June through August 2018. Mean±SD of continuously monitored environmental data are shown for periods of tidal submergence only (tidal height >0.3m), collected at two deployment locations within each bay

|  | **Fidalgo Bay** | **Port Gamble Bay** | **Skokomish River Delta** | **Case Inlet** |
| --- | --- | --- | --- | --- |
| **Temperature (°C)** | 15.4±1.5 | 15.0±1.0 | 16.2±2.7 | 16.8±1.7 |
| **DO (mg/L)** | 10.6±2.4 | 10.5±1.9 | 10.2±3.9 | 11.2±2.8 |
| **Salinity (PSU)** | 28.5±3.9 | 31.9±2.0 | 29.6±1.3 | 24.6±1.7 |
| **pH<sub>T</sub>** | 8.07±0.15 | 7.86±0.17 | 8.01±0.20 | 8.01±0.16 |
| **Chlorophyll (µg/L)** | 2.27±4.09 | 2.25±1.45 | 5.72±15.36 | 3.31±6.13 |

During both temperature and pCO_2_ treatments, all oysters were fed from a shared algae header tank daily with Shellfish Diet 1800® (300-500-mL, Reed Mariculture) diluted in ambient pCO_2_ seawater (200-L, Helm & Bourne, 2004), dosed continuously with metering pumps. Experimental, reservoir, and algae tanks were drained and cleaned, and oysters were monitored for mortality and rotated within the experimental system twice weekly.

### Adult reproductive development

A subset of oysters was sampled for gamete stage and dominant sex immediately before and after pCO_2_ treatments (Figure 2) to capture developmental differences among treatments. Puget Sound *O. lurida* reportedly enter reproductive quiescence and resorb residual gametes when temperatures are below 12.5°C (Hopkins 1936, 1937), however recent evidence of low-temperature brooding in Puget Sound (10.5°C, Barber *et al*. 2016) suggests that reproductive activity may occur during warm winters. Therefore, gonad tissue was sampled to estimate the following: 1) whether residual gametes were resorbed or developed during winter treatments; 2) whether temperature and pCO_2_ influenced winter activity; 3) if male and female gametes responded similarly; and 4) if gonad responses correspond with fecundity. Prior to pCO_2_ exposure, 15 oysters were sampled from O-1, O-2, and F cohorts, and 9 from the D cohort. After pCO_2_ exposure, 9, 6, and 15 oysters were sampled from each treatment for O-1/F, D, and O-2 cohorts, respectively (distributed equally among replicates tanks). Whole visceral mass was excised and preserved in histology cassettes using the PAXgene Tissue FIX System, then processed for gonad analysis by Diagnostic Pathology Medical Group, Inc. (Sacramento, CA).

Adult gonad samples were assigned sex and stage using designations adapted from (da Silva, Fuentes, & Villalba, 2009) (Appendix S1: Tables S2 & S3). Sex was assigned as indeterminate (I), male (M), hermaphroditic primarily-male (HPM), hermaphroditic (H), hermaphroditic primarily-female (HPF), and female (F). Gonad sex was collapsed into simplified male and female designations for statistical analyses (hermaphroditic-primarily male = male, hermaphroditic-primarily female = female). For stage assignment, male and female gametes were assigned separately due to the high frequency of hermaphroditism (50.8%). Dominant gonad stage was then assigned based on the sex assignment. The da Silva gonad stages were applied for early gametogenesis (stage 1), advanced (stage 2), and ripe (stage 3). Departures from da Silva’s stage 0, stage 4 (partially spawned), and stage 5 (fully spawned/resorbing) were as follows: stage 0 in this study represents empty follicles, or no presence of male or female gonad tissue; stage 4 represents both spawned and resorbing gonad; this method did not include a separate stage 5, due to the very high frequency of residual gametes, and no distinct partially spawned oysters (for gonad images see Appendix S1: Fig. S2 and Spencer *et al*. 2019).

Treatment effects on gonad tissue were assessed for all cohorts combined in 4 gonad metrics: 1) gonad stage of dominant sex, 2) male gonad tissue when present, 3) female gonad tissue when present, and 4) gonad sex-collapsed (Chi-square test of independence). To assess the effects of elevated winter temperature alone, gonad metrics were compared between 6°C and 10°C treatments prior to pCO_2_ treatment. To determine the effect of pCO_2_ exposure, gonad metrics were compared between ambient and high pCO_2_ after 52 days in pCO_2_ treatments, including temperature interaction effects. To estimate whether gonad changed during pCO_2_ treatment, metrics were compared before and after ambient and high pCO_2_ treatments, including temperature interaction effects. P-values were estimated using Monte-Carlo simulations with 1,000 permutations, and corrected using the Benjamini & Hochberg method and *α*=0.05 (Benjamini & Hochberg, 1995).

### Larval production

Following pCO_2_ exposure, adult oysters were spawned to assess impacts of winter treatment on larval production timing and magnitude. Beginning on April 11, 2017 (Figure 2), oysters were reproductively conditioned by raising temperatures gradually (∼1°C/day) to 18.1 ± 0.1°C and fed live algae cocktail at 66,000 ± 12,000 cells/mL. Oysters spawned in the hatchery for 90 days volitionally, i.e. naturally releasing gametes without chemical or physical manipulation. Six spawning tanks were used for each temperature x pCO_2_ treatment: 6°C-high pCO_2_, 6°C-ambient pCO_2_, 10°C-high pCO_2_, and 10°C-ambient pCO_2_. Within the six tanks per treatment, two spawning tanks contained the F cohort (14-17 oysters), two tanks the O-1 cohort (14-17 oysters), one tank the D cohort (9-16 oysters), and one tank the O-2 cohort (111-126 oysters). More O-2 oysters were used due to their small size. Olympia oysters release sperm, but have internal fertilization and release veliger larvae following a ∼2 week brooding period (Coe, 1931; Hopkins, 1937). Therefore, production was assessed by collecting veliger larvae upon maternal release. Spawning tank outflow was collected in 7.5-L buckets using 100 µm screens made from 15.25 cm polyvinyl chloride rings and 100 µm nylon mesh.

Larval collection was assessed for differences in spawn timing and fecundity. Larvae, first observed on May 11, 2017 (Figure 2), were collected from each spawning tank every one or two days for 60 days. The number of larvae released was estimated by counting and averaging triplicate subsamples of larvae homogenized in seawater. The following summary statistics were compared between temperature x pCO_2_ treatments: total larvae released across the 90-day period, average number of larvae collected on a daily basis (excluding days where no larvae were released), maximum larvae released in one day, date of first release, date of maximum release, and number of substantial release days (greater than 10,000 larvae). The total and daily release values were normalized by the number of broodstock * average broodstock height (cm), which can impact fecundity. Distributions were assessed using `qqp` in the `car` package for R (Fox & Weisberg, 2011), and log-transformed to meet normal distribution assumptions, if necessary.

Differences between treatments were assessed using linear regression and Three-Way ANOVA (cohort was included as a covariate) with backwards deletion to determine the most parsimonious models. Tukey Honest Significant Differences were obtained using `TukeyHSD` to assess pairwise comparisons (R Core Team, 2016). Dates of peak larval release were also estimated for each pCO_2_ x temperature treatment by smoothing using locally weighted regression, with `geom_smooth` in the ggplot package (Wickham, 2017), with span=0.3 and degree=1.

### Offspring survival in a natural setting

To assess potential carryover effects of parental pCO_2_ exposure, offspring from parents in 6°C-ambient pCO_2_ and 6°C-high pCO_2_ treatments were reared then deployed in the natural environment. To focus on the effect of parental pCO_2_ exposure, only offspring from 6°C parents were tested in the field (Figure 2). Larvae were collected between May 19 and June 22, 2017, separated by parental pCO_2_ exposure and cohort, and reared in common conditions for approximately 1 year (Figure 2; for rearing methods see Appendix S1: Section S6). On June 12, 2018 the juveniles were placed in four bays in Puget Sound — Fidalgo Bay, Port Gamble Bay, Skokomish River Delta, and Case Inlet — with two sites per bay, for a total of eight locations (Figure 1). Autonomous sensors collected continuous water quality data at each location for pH, salinity (via conductivity), dissolved oxygen, temperature, and chlorophyll. For the F/D and O-1/O-2 cohorts, respectively, 30 and 10 oysters were placed at each location. Initial shell height and group weight were measured, then oysters were enclosed in mesh pouches and affixed inside shellfish bags to exclude predators. At the end of three months, survival, shell height and group weight were measured for live oysters.

Juvenile oyster survival was compared among bays and parental pCO_2_ exposure with a binomial generalized linear mixed model (glmm) using `glmer` from the `lme4` package (vs. 1.1-19). Chi-square tests compared survival differences among factors using the `car` package `Anova` function (Fox & Weisberg, 2011). Mean shell growth was determined by subtracting pre-deployment mean height from post-deployment mean height (not including dead oysters). Both mean shell growth and mass change were compared among factors using ANOVA and F-statistics to test differences by bay, parental pCO_2_, and cohort.

Make and model details for instruments used during treatments and field deployments are available in the Appendix S1: Section S2. All data analysis was performed in R version 3.3.1 using the RStudio interface (R Core Team, 2016). Code for statistical analyses can be found in the associated Github repository (Spencer *et al.,* 2019).

## Results

### Adult reproductive development

After 60 days in temperature treatments (6.1±0.2°C and 10.2±0.5°C), gonad stage of the dominant sex differed significantly between temperatures (Table 2). The 10°C oysters had more instances of advanced gametogenesis (stage 2), and fewer that were resorbing/spawned (stage 4) (Figure 3). This difference was influenced strongly by more advanced male gametes in 10°C oysters, but there were no differences in female gamete stages. No differences in sex ratio were observed between temperature treatments (Figure 4).

**Table 2:** Gonad stage and sex comparisons among treatments. Gonad was sampled after temperature treatment but before pCO_2_ (6°C Pre and 10°C Pre, n=54), and after pCO_2_ treatment (Amb=841±85 µatm, n=39; High= 3045±488 µatm, n=39). Pearson’s chi-square statistics are shown with p-adj in parentheses for gonad sex, stage of the dominant sex, male gametes when present, and female gametes when present. Cells with * and in bold indicate significant differences between comparison; blank cells=not tested; % of mature = % of sampled oysters that contained stage 3 male or female gametes, per treatment.

| Temperature | pCO<sub>2</sub> | 6°C |  |  | 10°C |  |  | 6°C |  |  | 10°C |  |  |
| --- | --- | --- | --- | --- | --- | --- | --- | --- | --- | --- | --- | --- | --- |
|  |  | Pre | Amb | High | Pre | Amb | High | Pre | Amb | High | Pre | Amb | High |
| 6°C | Pre | - |  |  |  |  |  | - |  |  |  |  |  |
|  | Amb | 0.8<br>(0.93) | - |  | *Sex Ratio* |  |  | *16.5<br>(0.013) | - | *Stage of the dominant sex* |  |  |  |
|  | High | 4.6<br>(0.34) | 5.4<br>(0.29) | - |  |  |  | 4.6<br>(0.48) | 9.7<br>(0.090) | - |  |  |  |
| 10°C | Pre | 5.9<br>(0.26) |  |  | - |  |  | *15.8<br>(0.017) |  |  | - |  |  |
|  | Amb |  |  |  | 6.8<br>(0.18) | - |  |  |  |  | *12.7<br>(0.038) | - |  |
|  | High |  | 5.3<br>(0.29) |  | 3.8<br>(0.46) | 0.6<br>(0.94) | - |  | 2.8<br>(0.78) |  | 5.2<br>(0.44) | *12.5<br>(0.038) | - |
| 6°C | Pre | - |  |  |  |  |  | - |  |  |  |  |  |
|  | Amb | *24.2<br>(1.6e-3) | - |  | *Male gametes* |  |  | 6.3<br>(0.18) | - | *Female gametes* |  |  |  |
|  | High | *15.2<br>(0.013) | 9.0<br>(0.071) | - |  |  |  | 3.6<br>(0.47) | 4.4<br>(0.36) | - |  |  |  |
| 10°C | Pre | *31.1<br>(1.6e-3) |  |  | - |  |  | 2.1<br>(0.78) |  |  | - |  |  |
|  | Amb |  |  |  | *11.2<br>(0.038) | - |  |  |  |  | 4.2<br>(0.26) | - |  |
|  | High |  | 1.7<br>(0.78) |  | 0.6<br>(0.95) | 9.5<br>(0.084) | - |  | 0.8<br>(0.9) |  | 5.5<br>(0.17) | 0.15<br>(1.0) | - |
| % mature |  | 30% | 28% | 15% | 19% | 33% | 21% | 2% | 15% | 8% | 6% | 18% | 21% |

**Figure 3:**
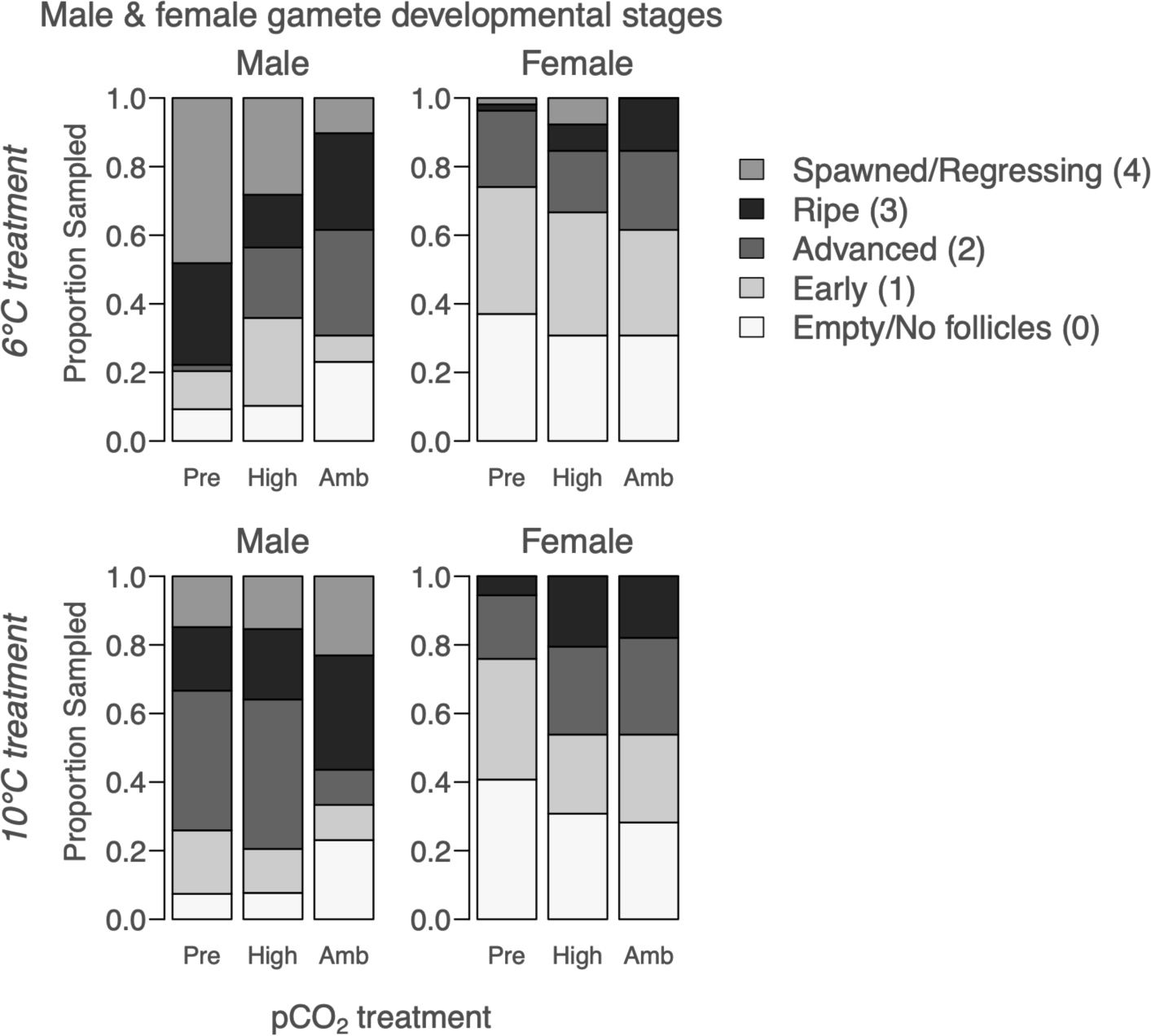
Gonad developmental stages for male and female gametes, after 60-days in temperature treatments but before pCO_2_ treatments (“Pre”, n=54) and after 52 days in high pCO_2_ (3045±488 µatm, n=39) and ambient pCO_2_ (841±85 µatm, n=39), which indicates that sperm development was influenced by elevated winter temperature (more advanced) and high pCO_2_ (less advanced, 10°C treatment only), but oocyte development was not. All oysters were assigned both male & female stages; if no oocytes were present, for example, that oyster was designated as female stage 0.

**Figure 4:**
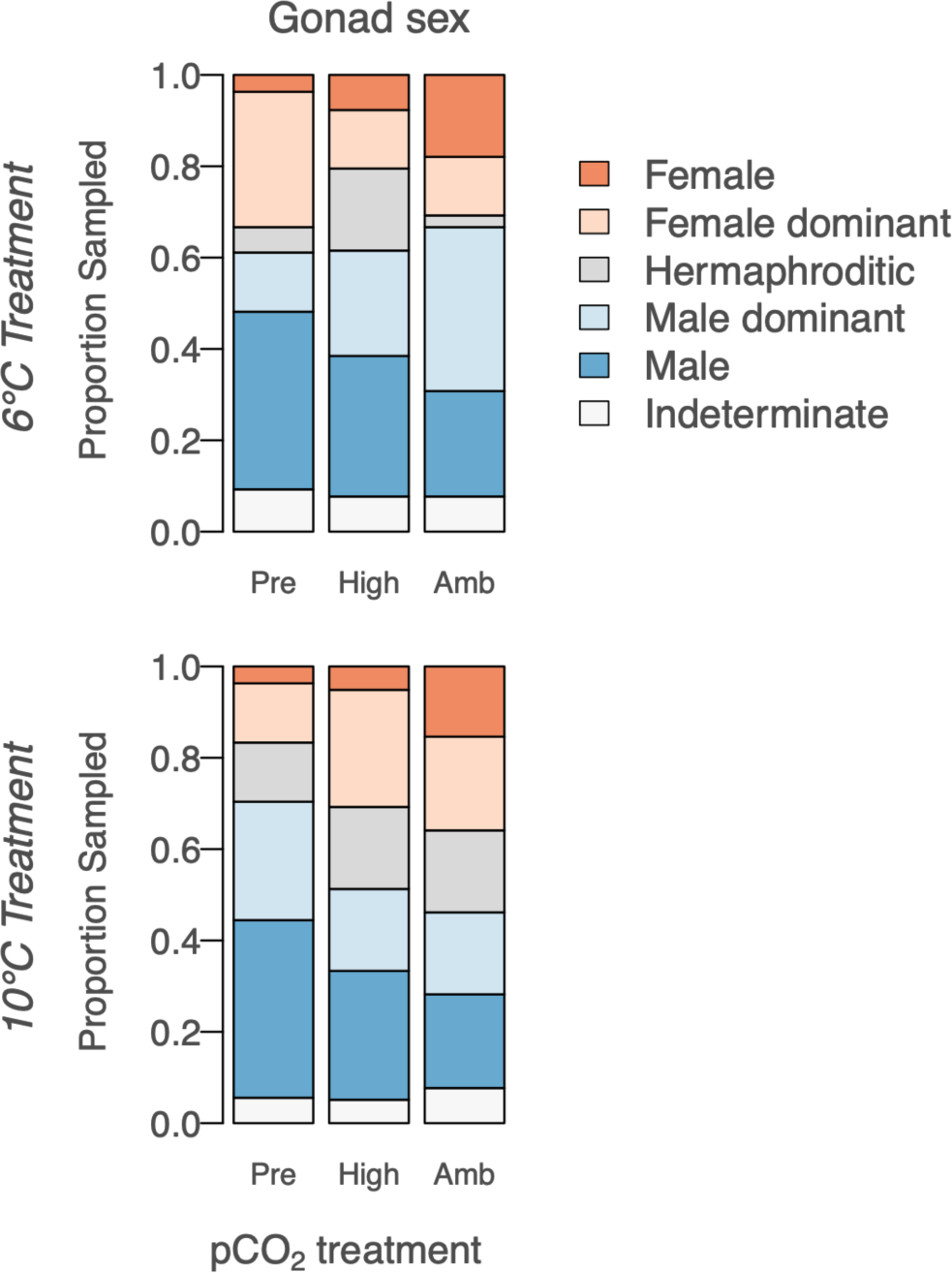
Gonad sex, after 60-days in temperature treatments but before pCO_2_ treatments (“Pre”, n=54) and after 52 days in high pCO_2_ (3045±488 µatm, n=39) and ambient pCO_2_ (841±85 µatm, n=39). Winter conditions did not significantly influence gonad sex ratios.

After 52 days in pCO_2_ treatments, gonad stage of the dominant sex differed significantly between ambient and high pCO_2_ in the oysters previously held in 10°C (Table 2). More mature gametes (stage 3) were found in 10°C-ambient pCO_2_ (49%) compared to 10°C-high pCO_2_ (33%). This difference was strongly influenced by oysters that were predominantly male, as male gamete stage tended to differ between pCO_2_ treatment, but female gamete stage did not (Table 2, Figure 3). In 6°C-treated oysters, there were no pCO_2_ effects on gonad stage of the dominant sex, male gamete stage, or female gamete stage. No gonad stage or sex ratio differences were detected among oysters from 10°C-high pCO_2_ (combined stressors) and 6°C-ambient pCO_2_ (no stressors). Gonad sex did not differ significantly among treatments, however oysters tended to contain fewer male-only and more female-only gonad tissues in the riper, ambient pCO_2_-treated groups than male-only tissues (Figure 4).

Compared to oysters before pCO_2_ exposure, those exposed to high pCO_2_ did not differ in gonad sex, stage of the dominant sex, or female gamete stage. Male gametes in the 6°C treated oysters developed while in the high pCO_2_ exposure, but there was no change in the 10°C treated oysters. Oysters held in ambient pCO_2_ had significantly more advanced gonad compared to before CO_2_ exposure regardless of temperature, again influenced strongly by changes in male gamete stage (Table 2).

No sampled oysters contained brooded embryos or larvae. Gonad data and patterns within cohorts is reported in Appendix S1: Figures S3, S4, and Table S4.

### Larval production

Adults exposed to 10°C produced more larvae on a daily basis (excluding days where no larvae were released) than those exposed to 6°C in ambient pCO_2_-exposed oysters (p=0.040), but not in high pCO_2_-exposed oysters (p=0.66) (Figure 6, pCO_2_:temperature interaction: (F(2,8)=5.1, p=0.037). Total larvae released over the 90-day spawning period tended to differ by treatment, but not significantly (temperature:pCO_2_ interaction (F(2,8)=4.0, p=0.063). Temperature and pCO_2_ as single factors did not affect total larvae released or daily averages.

The date of first larval release differed by temperature regardless of pCO_2_ (Figures 5 & 6, F(1,8)=11.9, p=0.0087), and pCO_2_ had no effect on timing (not retained in model). Onset was on average 5.2 days earlier in the 10°C treatment. Timing of peak larval release also differed by temperature treatment regardless of pCO_2_ (Figure 6, F(3,19)=6.7, p=0.018), occurring on average 8.3 days earlier in 10°C oysters. The 10°C treated oysters produced more large pulses of larvae, on average 2 additional days, than 6°C (F(1,8=7.25, p=0.027).

**Figure 5:**
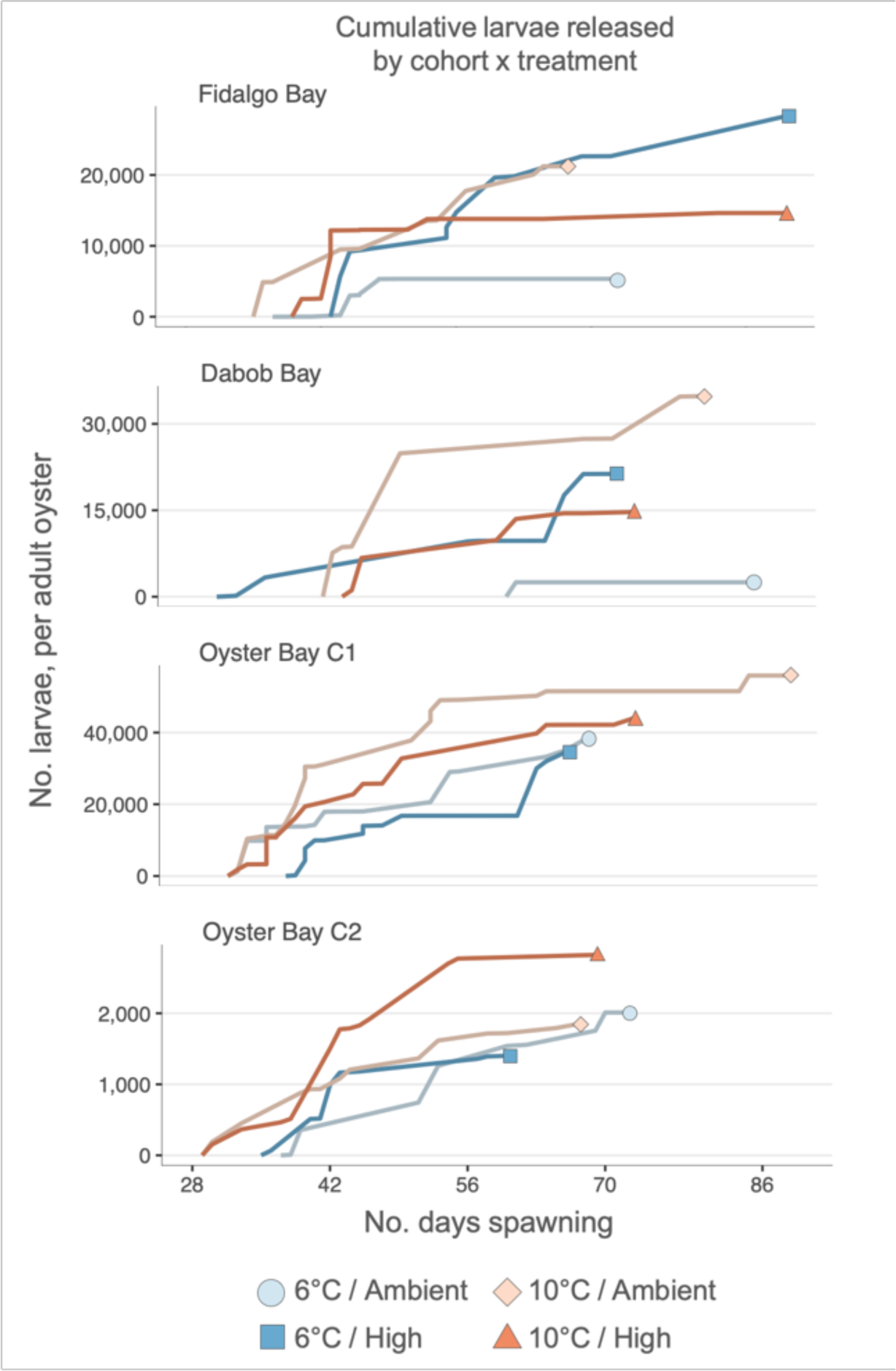
Cumulative larvae released over 90 days of continuous volitional spawning under hatchery conditions, normalized by the number of adult oysters. Each of the four panels represent a cohort, and lines are color coded by winter temperature and pCO_2_ treatments, where ambient pCO_2_ = 841 µatm (7.8 pH), and high pCO_2_ = 3045 µatm (7.31). Reproductive conditioning and spawning occurred at 18°C, in ambient pCO_2_, and with live algae at a density of 66,000 ± 12,000 cells/mL.

**Figure 6:**
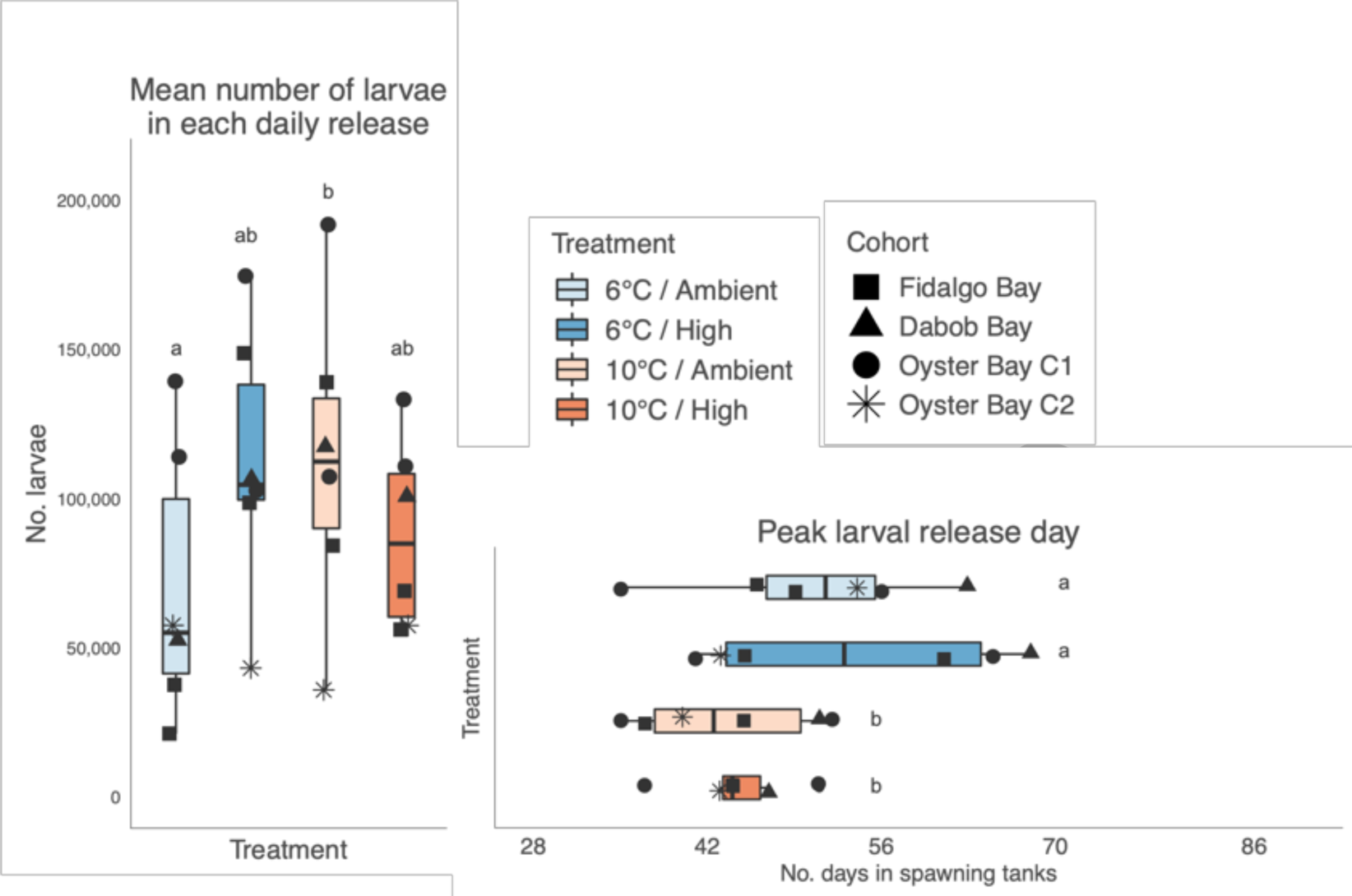
Left: average number of larvae collected on a daily basis (excluding days where no larvae were released). Daily pulses of larvae were larger in 10°C than 6°C, but only in oysters exposed to ambient pCO_2_. For statistical analysis, data was normalized by number of oysters * average oyster height (cm) (data shown is not normalized). Right: number of spawning days until larval release peaked; peak release occurred on average 8.3 days earlier in 10°C treated oysters. Letters (a, ab, b) indicate differences among treatments. Boxes contain values lying within the interquartile range (IQR), with medians indicated by lines in the middle of boxes. Whiskers extend to the largest value no greater than 1.5*IQR.

In total, 18.5 million larvae were collected from 767 oysters. Total larvae produced by each treatment was 3.1M, 4.8M, 5.9M, and 4.5M for 6°C-ambient pCO_2_, 6°C-high pCO_2_, 10°C-ambient pCO_2_, and 10°C-high pCO_2_, respectively. Based on reports of approximately 215,000 larvae produced per adult *O. lurida* of shell height 35 mm (Hopkins, 1936), the number of oysters that spawned as female in this study was approximately 86, with 14.3, 22.5, 27.6, and 21.0 from the 6°C-ambient pCO_2_, 6°C-high pCO_2_, 10°C-ambient pCO_2_, and 10°C-high pCO_2_ treatments, respectively. This estimate is likely low across all treatments, due to the smaller D and O-2 cohorts (mean length in F, D, O-1 and O-2 was 35.7 mm, 29.8 mm, 35.7 mm, and 20.0 mm, respectively), therefore the total number of oysters that spawned as female and released larvae is likely higher than 86.

Larval production and timing data, including differences among cohorts, are included in Appendix S1: Section S5 and Table S5.

### Offspring survival in a natural setting

Juvenile survival after three months in the field was on average 15% higher in cohorts from high pCO_2_ exposed parents than from ambient pCO_2_ parents (44±37%, and 29±27%, respectively, *χ*^2^=10.6, p=0.0011). The influence of parental pCO_2_ on survival varied by bay (bay:parental pCO_2_ interaction *χ*^2^=15.3, p=1.6e-3), and by cohort (cohort:parental pCO_2_ interaction *χ*^2^=23.5, p=3.2e-5) (Table 3).

**Table 3:**
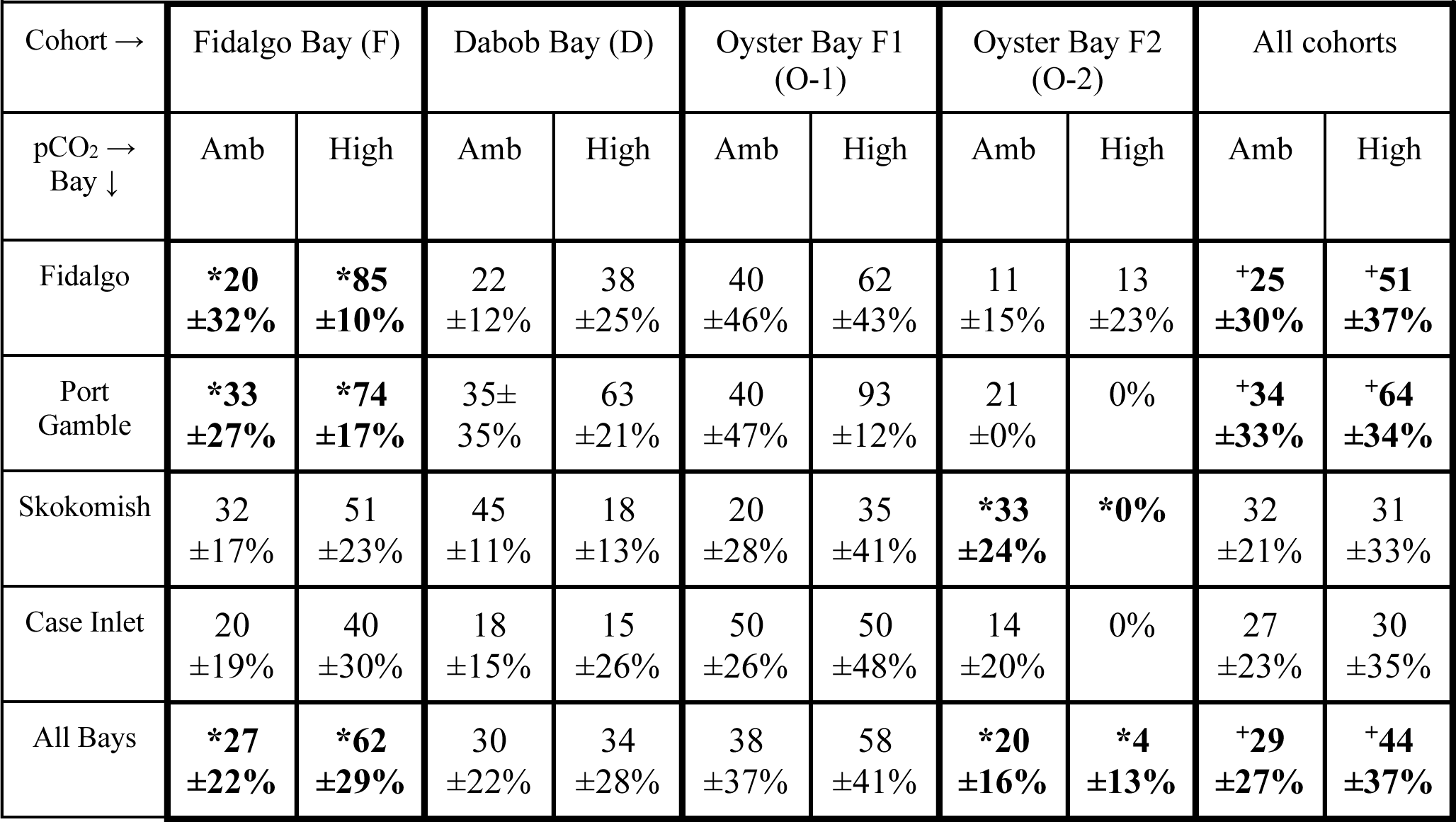
Offspring survival in the field. 1-year old juveniles were deployed for 3 months in four bays in Puget Sound, Washington, in 2 sites per bay. Percent survival ± SD is shown by cohort x bay x parental pCO_2_ treatment (Amb=841±85 µatm, High= 3045±488 µatm). Only offspring from 6°C-treated adults were deployed. Significant survival differences were detected between parental pCO_2_ treatment within the Fidalgo Bay and Oyster Bay F2 cohorts (*), and across all cohorts (+).

Survival in offspring from high pCO_2_ parents was higher in the Fidalgo Bay and Port Gamble Bay locations (*χ*^2^=17.7, p= 2.6e-5; *χ*2=10.0, p=1.6e-3, respectively), but this was not the case in Skokomish River Delta or Case Inlet. Survival in the F cohort was 38% higher in oyster from pCO_2_ parents than those from ambient pCO_2_ parents across all deployment bays (*χ*^2^=28.1, p=4.6e-7), and within the Fidalgo Bay location (*χ*^2^=17.6, p-adj=0.0001). Survival in the D and O-1 cohorts did not differ significantly between parental pCO_2_ across all bays (D: *χ*^2^=0.4, p=1, O-1: *χ*^2^=2.5, p=0.44), or within individual bays. More O-2 juveniles with ambient pCO_2_ parents survived across all bays (*χ*^2^=9.1, p=0.010), and within the Skokomish River Delta (*χ*^2^=8.9, p=0.011).

Without considering parental pCO_2_, more oysters survived in Port Gamble Bay (mean 49±36%) and Fidalgo Bay (39±36%) than in Case Inlet (mean 29±29%, p=0.012 & p=0.037, respectively) (bay factor, *χ*^2^=18.5, p=3.4e-4). Survival at Skokomish River Delta did not differ significantly from other locations (32±27%). No interaction between cohort and bay was detected (*χ*^2^=9.8, p=0.37) (Figure 7, Table 3).

**Figure 7:**
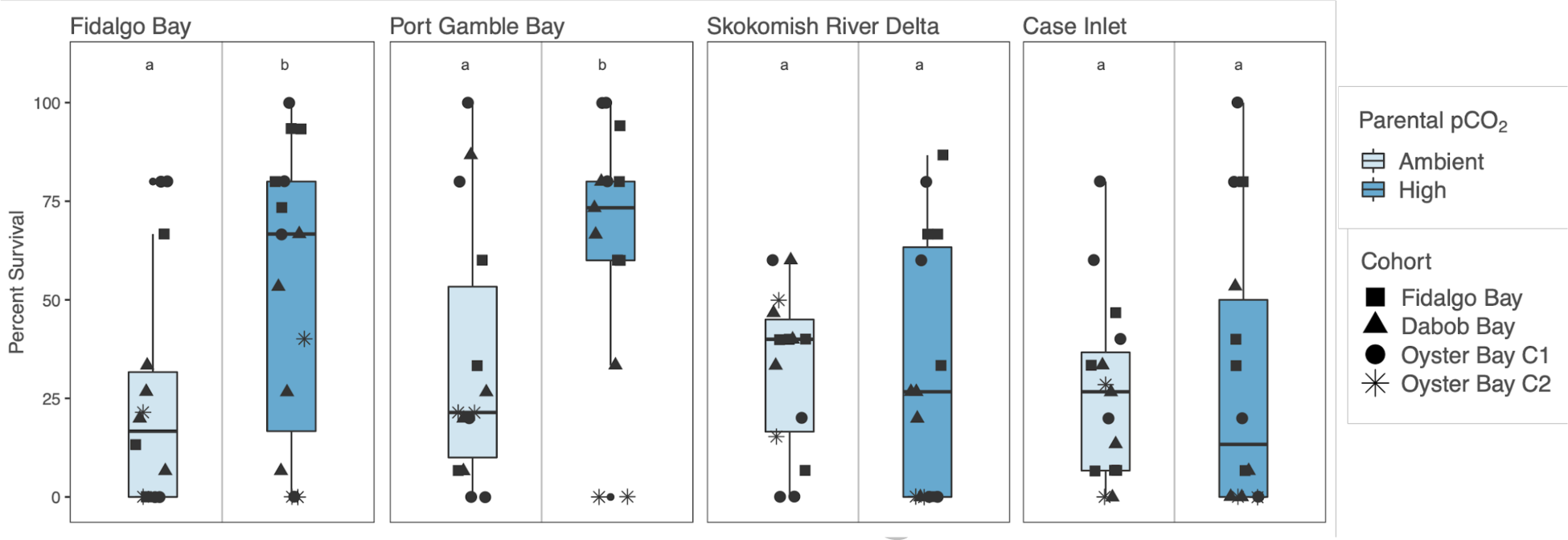
Percent survival of juvenile offspring in the field. The four panels each represent survival in one bay (Fidalgo Bay, Port Gamble Bay, Skokomish River Delta, Case Inlet). Within each panel, boxplots are separated by parental pCO_2_ exposure (Ambient=841 µatm, High=3045 µatm). Points indicate % survival in each deployment pouch, and symbols indicate cohort (Fidalgo Bay, Dabob Bay, Oyster Bay Cohort 1, and Oyster Bay Cohort 2). Letters (a, b) indicate survival differences among parental pCO_2_ exposure within each bay. Boxes contain values lying within the interquartile range (IQR), with median survival indicated by lines in the middle of boxes. Whiskers extend to the largest value no greater than 1.5*IQR.

Shell length was not affected by bay, cohort or parental pCO_2_. The mass per oyster (compared to before deployment) differed by cohort (F(3,76)=15.9, p=4.0e-8), due to Dabob Bay cohort growing less than the other three cohorts (Δ g/oyster: D=0.5, F=1.2, O-1=1.6, & O-2=1.0). Mass change also differed by bay (F(3,76)=4.8, p=3.9e-3) due to less growth in oysters placed at Fidalgo Bay than in Port Gamble Bay and Case Inlet (Δ g/oyster: FB=0.7, PGB=1.0, CI=1.1, SK=0.8) (Appendix S1: Figure S5).

## Discussion

Ocean acidification and ocean warming potentially threaten marine organisms, particularly ectothermic calcifiers (Hoffman *et al*. 2010). An organism’s genotype, complete environmental history, and the timing and magnitude of environmental perturbations may all determine its fitness in future ocean conditions. To begin teasing apart these complex factors in the Olympia oyster, this study examined four adult cohorts with distinct genetic structure but known, shared histories. Elevated winter temperature resulted in increased gonad development, which corresponded with earlier and more frequent larval release (on average 5.2 days earlier, 2 additional days). High pCO_2_ exposure negatively influenced gonad maturation state, but did not affect subsequent fecundity. Offspring from parents exposed to elevated pCO_2_ had higher overall survival upon deployment. Differences in juvenile survival among bays and cohorts indicate that carryover effects are dependent upon the environment and genotype, and reinforce the importance of using multiple sources of test organisms in stress-response studies.

### Reproduction

We expected elevated winter temperature to reduce fecundity, based on predictions that changes to reproductive quiescence and metabolism would be deleterious to spring reproduction. Counter to this prediction, warm winter temperature positively affected larval production. Oysters in elevated temperature contained more developed male gametes after treatment, and subsequently began releasing larvae earlier and produced more larvae per day compared to cold-treated oysters. We find no evidence that cold winters are critical for spring reproduction, but rather elevated winter temperature may elongate the *O. lurida* spawning season. In comparison, a 29-year dataset of *M. balthica* reproduction showed that as winter temperature increased, spring spawning began earlier and fecundity declined (Philippart *et al*., 2003). However, the present study was conducted in a hatchery setting, with ample phytoplankton, and did result in a temperature shift during spawning. In the wild numerous additional abiotic and biotic factors will contribute to *O. lurida* fitness, and warmer winters may result in earlier and longer reproductive seasons only if nutritional requirements are met. Whether larvae released earlier in the spring can survive to recruitment will greatly depend on many factors including food availability and predation. Those modeling larval recruitment (*e.g.* Kimbro, White & Grosholz, 2019; Wasson *et al*., 2016) should consider including winter temperature as a factor influencing spatiotemporal recruitment patterns.

We predicted that high pCO_2_ exposure would redirect energy away from storage to maintenance processes, resulting in delayed gametogenesis and poor fecundity in the spring. After exposure to 3045 µatm pCO_2_ (pH 7.31), fewer oysters contained ripe or advanced male gonad tissue than in ambient pCO_2_, signaling reduced spermatogenic activity. Female gonad, sex ratios, and subsequent fecundity were not affected by sole exposure to high pCO_2_. Similar impacts on gametogenesis during exposure were observed in the Sydney rock (*S. glomerata*) and Eastern (*C. virginica*) oysters, but with varying pCO_2_ thresholds. Parker *et al*. (2018) found *S. glomerata* gametogenesis to slow in 856 µatm (pH 7.91), and Boulais *et al*. (2017) found normal rates at 2260 µatm (pH 7.5), delay at 5584 µatm (pH 7.1), and full inhibition at 18480 µatm (pH 6.9) in *C. virginica*. Together, these studies indicate that high pCO_2_ slows the rate of gametogenesis, but the level at which pCO_2_ affects gametogenesis appears species-specific, and likely reflective of variable physiological mechanisms and reproductive strategies.

The combined effects of sequential elevated temperature and pCO_2_ treatments did not act synergistically to delay gonad development, but instead resulted in oysters with gonad stage and fecundity no different from the untreated oysters. Similarly, combined simultaneous temperature and high pCO_2_ exposures did not affect *S. glomerata* fecundity (Parker *et al*., 2018). We did detect a pCO_2_ dependent effect of temperature on the average number of larvae released per day. Oysters that had previously been exposed to 10°C produced more larvae than 6°C, but only after ambient pCO_2_ exposure, which may reflect a general reproductive arrest that occurs when exposed to high pCO_2_. Despite experimental differences (*e.g.* sequential vs. simultaneous exposures) which can influence outcomes (Bible *et al*. 2017), both Parker *et al*. (2018) and the present study indicate that high pCO_2_ slows gametogenesis, elevated temperature accelerates it, and these two environmental drivers act antagonistically on gonad development if occurring in the same reproductive season. An important factor not included in either study is ecologically relevant variability. Temperature and pCO_2_ oscillations, driven by tides and diurnal photosynthesis, could offer daily refuge or expose oysters to dynamic changes, altering how combined stressors interact (Cheng *et al*. 2015).

In contrast to prior studies, temperature and pCO_2_ did not impact *O. lurida* sex ratios, whereas in high pCO_2_ *C. virginica* skewed male (Boulais *et al.,* 2017), and *S. glomerata* skewed female (Parker *et al*., 2018). This observation may be explained by very low incidence of total reproductive inactivity in our *O. lurida* cohorts — only four out of the 108 oysters that were sampled prior to pCO_2_ treatment contained empty follicles — and thus sex ratios may be different if pCO_2_ exposure occurs earlier in life during initial sex differentiation. Furthermore, high pCO_2_ exposure only occurred in winter, prior to spawning. If high pCO_2_ persists during oocyte maturation and spawning, *O. lurida* fecundity may be reduced similar to *C. virginica* and *S. glomerata.* Future research should examine *O. lurida* sexual development during the initial switch from male to female, which can occur the first winter after settlement (Moore *et al*., 2016), and across a range of pCO_2_ to determine conditions in which gametogenesis and sex determination are affected.

### Offspring

Abiotic parental stressors can be beneficial, neutral, or detrimental to offspring viability (Donelson *et al.,* 2018). We explored carryover effects of adult exposure to winter pCO_2_ on offspring by testing survival in the field. Offspring with high pCO_2_ parental histories performed better in two of four locations, Fidalgo Bay and Port Gamble Bay. Carryover effects of parental high pCO_2_ exposure may therefore be neutral, or beneficial, to offspring depending on the environmental conditions. Port Gamble Bay and Fidalgo Bay are more influenced by oceanic waters, which could explain cooler observed temperatures. These locations are also typically less stratified than the Skokomish River Delta and Case Inlet. In Port Gamble Bay, where pCO_2_ parental history most significantly correlated with offspring survival across cohorts, mean pH was considerably lower than the other deployment locations (-0.17 pH units), and mean salinity was higher (+3.8 PSU). Given the experimental design we are able to clearly demonstrate that manifestation of carryover effects in Olympia oysters is dependent on environmental conditions. Specifically, there is a greater likelihood of beneficial carryover effects when parents are exposed to stressful conditions. Overall, carryover effects of parental pCO_2_ treatment were positive, however negative effects were observed in the O-2 cohort. This discrepancy could relate to unique O-2 juvenile characteristics, as they were bred from siblings, and were 3rd-generation hatchery produced. The complex interactions among parental exposure, bay, and cohort indicate that offspring viability is influenced by ancestral environment history, environmental conditions, and genotype.

Our results contrast with a similar study that exposed *C. gigas* oysters to high pCO_2_during the winter, and found fewer hatched larvae 18 hours post-fertilization from exposed females, with no discernable paternal effect (Venkataraman, Spencer & Roberts, 2019). Hatch rate was not directly measured in this study due to the *O. lurida* brooding behavior; however, no difference in daily and total larvae released suggest that hatch rate was unaffected by pCO_2_. The different responses seen in Venkataraman, Spencer & Roberts (2019) and the present study may reflect variability among species and spawning method. *C. gigas* gametes were collected artificially by stripping gonad, whereas *O*. *lurida* late-stage veliger larvae were collected upon release from the brood chamber. For instance, volitionally-spawned gamete quality and fertilization rates could vary between the natural versus artificial settings to influence larval viability. Larval brooding may also be a mechanism by which sensitive larvae are acclimatized to stressors, as the *O. lurida* brood chamber pH and dissolved oxygen can be significantly lower than the environment (Gray *et al*., *in press*).

Beneficial parental carryover may also be linked to the male-specific gonad effects, and the conditions in which the adult oysters were held. During high pCO_2_ exposure, oocyte stage and prevalence did not change, which indicates that oogenesis did not occur. Negative intergenerational carryover effects are commonly linked to variation in oocyte quality, which can be affected by the maternal environment during oogenesis (Utting & Millican, 1997). In the Chilean flat oyster (*Ostrea chilensis*), for instance, egg size and lipid content positively correlate with juvenile growth and survival (Wilson, Chaparro, & Thompson, 1996). If high pCO_2_ exposure were to coincide with oocyte proliferation and growth, *O. lurida* egg quality and larval viability could be compromised. In contrast, male gonad stage advanced significantly during pCO_2_ exposure. Intergenerational and transgenerational carryover effects are increasingly linked to the paternal environment in other taxa, such as inheritance of epigenetic changes to the male germ line (Rodgers, Morgan, Bronson, Revello, & Bale, 2013; Anway, 2005; Soubry, Hoyo, Jirtle, & Murphy, 2014). Positive carryover effects of environmental stressors observed in this and other marine invertebrate taxa may be due to paternal epigenetic effects, but this link has not yet been observed.

## Conclusion

This study clearly demonstrates that exposure to elevated winter temperature and altered carbonate chemistry impacts reproduction and offspring viability in the Olympia oyster. Furthermore, we report the first observations of intergenerational plasticity in an *Ostrea* species, that is dependent on offspring environmental conditions and population. The observed context-dependent carryover effects could have a substantial impact on species resilience. Combined with previous reports of resilience to environmental stressors (Waldbusser *et al* 2016; Cheng *et al*. 2017) and intraspecific variability (Bible, Evans & Sanford, 2019; Maynard, Bible, Pespeni, Sanford, & Evans, 2018; Silliman, Bowyer, & Roberts, 2018; Heare, Blake, Davis, Vadopalas, & Roberts, 2017), the Olympia oyster may be more capable than other marine bivalve species to withstand and adapt to unprecedented ocean change. Furthermore, conserving and restoring *O. lurida* in a variety of settings — including hypoxic, warmer, and less alkaline areas — could increase the probability that future populations are equipped for challenging conditions through selection or intergenerational carryover.

As temperatures rise and ocean acidification progresses, there may be profound and unexpected seasonal changes across marine taxa. Accurate predictions will need to consider parental carryover effects, as they can impart neutral, beneficial, or detrimental characteristics to offspring, which depend on complex interactions among parental exposure timing, reproductive strategies, species plasticity, and standing genetic structure. With these considerations, future biological response studies need to be aware of three possible factors influencing results: 1) source population; 2) environmental history (within-lifetime carryover effects); and 3) ancestral environmental history (inter- and transgenerational carryover effects). Controlling for, or at minimum recognizing and recording these factors, will provide important context for those predicting ecosystem response to environmental change.

## Supporting information

Suppplemental materials

## Acknowledgements

Our gratitude to the following people who assisted with this project: Grace Crandall, Kaitlyn Mitchell, Olivia Smith, Megan Hintz, Rhonda Elliott, Lindsay Alma, Duncan Greeley, Beyer and Jackson Roberts, and Ian Davidson helped with oyster husbandry and sampling; Alice Helker advised on husbandry and larval rearing system engineering; Emily Kunselman helped manage the field deployment; Sam White and Hollie Putnam contributed to the carbonate chemistry analysis; Katherine Silliman and Jake Heare produced (and saved) the experimental oysters; the NOAA Manchester Research Center and Puget Sound Restoration Fund provided facilities and materials; committee members Jackie Padilla-Gamiño and Rick Goetz advised and supported this extended project. Thank you to David Kimbro and an anonymous reviewer for constructive comments on the manuscript.

This work was supported in part by the National Science Foundation Graduate Research Fellowship Program, the National Shellfisheries Association Melbourne R. Carriker Student Research Grant, Washington State Department of Natural Resources, and a grant from Washington Sea Grant, University of Washington, pursuant to the National Oceanic and Atmospheric Administration Award No. NA14OAR4170078; Project R/SFA-8. The views expressed herein are those of the author(s) and do not necessarily reflect the views of any funding agency.

