## Supplementary material for "Carryover effects of temperature and pCO_2_ across multiple Olympia oyster populations": Suppplemental materials

### Appendix S1

**Carryover effects of temperature and $pCO_2$ across multiple Olympia oyster populations.**
Spencer, L. H., Y. R. Venkataraman, R. Crim, S. Ryan, M. J. Horwith, & S. B. Roberts.

Data, R script analyses, gonad tissue images, and high-resolution supplementary figures are available in the associated GitHub repository: https://doi.org/10.6084/m9.figshare.8872646

#### Section S1: Recipe for Tris buffer (0.08 M, 28.0 PSU)

- 0.3603 mol of NaCl (J.T. Baker)
- 0.0106 mol of KCl (Fisher Scientific)
- 0.0293 mol MgSO4-(H2O)7 (Fisher Scientific)
- 0.0107 mol of CaCl2-2(H2O) (MP Biomedicals)
- 0.0401 HCl (J.T. Baker)
- 0.0799 mol of Tris base (Fisher Scientific)

Deionized water was added for a final volume of 1L

#### Section S2: Instruments used in experimental treatments in the hatchery

- Temperature treatment manipulation and monitoring:
  - Chilling experimental water: Teco Aquarium Chiller, Model TK-500
  - Continuous temperature measurements: Onset HOBO Water Temperature Data Loggers, Model U22-001
- **$pCO_2$ treatment manipulation and monitoring:**
  - pH monitoring: Durafet pH probes, Honeywell Model 51453503-505
  - $pCO_2$ injection control: Honeywell Dual Input Analytical Analyzer, Model 50003691-501
  - Temperature monitoring: HOBO Pendant Temperature Data Loggers, Model UA-002-64
- **Carbonate chemistry measurements:**
  - Temperature: Fisher Traceable Digital Thermometer, Model 15-077
  - Salinity: VWR Bench/Portable Conductivity Meter, Model 23226-505
  - pH: Mettler Toledo Combination pH Electrode, Model 11278-220
  - Total alkalinity: Mettler Toledo Excellence Titrator, Model T5 Rondolino
- Algal dosing: metering pump – Iwaki EZ Controller, Model EZCD1

**Instruments used in field trial for continuous environmental data collection:**
- pH: Honeywell Durafet II Electrode, in custom-built housing
- Salinity: via conductivity, Dataflow Systems Ltd. Odyssey Conductivity and Temperature Logger
- Dissolved Oxygen: Precision Measurement Engineering MiniDOT Logger
- Temperature: via dissolved oxygen probes
- Chlorophyll: Turner Designs Cyclops-7F Submersible Sensor with PME Cyclops-7 Data Loggers

**Table S1:** Carbonate chemistry parameters for three time points during the $pCO_2$ treatments, which are averages (± SE) from three replicate tanks per treatment. All parameters except for total alkalinity differed significantly between control/ambient (Amb.) and experimental/high (High.) tanks (One-way ANOVA). More details are available in Venkataraman, Spencer & Roberts, *2019.*

| Day | pH*** F(1,16) = 5838, p = 6.12e-22 | | Total Alkalinity (μmol/kg) F(1,16) = 1.38, p = 0.257 | | $pCO_2$ (μatm)*** F(1,16) = 235, p = 5.44e-11 | | DIC (μmol/kg)* F(1,16) = 7.12, p = 0.0168 | | $\Omega_{calcite}$*** F(1,16) = 529, p = 1.10e-13 | | $\Omega_{aragonite}$*** F(1,16) =527, p = 1.14e-13 | |
|---|---|---|---|---|---|---|---|---|---|---|---|---|
| | Amb. | High | Amb. | High | Amb. | High | Amb. | High. | Amb. | High | Amb. | High |
| 5 | 7.82 ± 0.004 | 7.33 ± 0.002 | 2307.41 ± 25.45 | 2332.36 ± 31.05 | 747.51 ± 13.94 | 2481.23 ± 29.83 | 2233.41 ± 25.29 | 2408.51 ± 31.76 | 1.86 ± 0.02 | 0.62 ± 0.01 | 1.16 ± 0.012 | 0.58 ± 0.007 |
| 33 | 7.81 ± 0.005 | 7.31 ± 0.004 | 2747.00 ± 21.13 | 2917.60 ± 18.36 | 912.22 ± 12.69 | 3309.52 ± 7.22 | 2664.57 ± 19.99 | 3020.99 ± 17.99 | 2.23 ± 0.03 | 0.77 ± 0.02 | 1.40 ± 0.020 | 0.48 ± 0.014 |
| 48 | 7.82 ± 0.015 | 7.29 ± 0.004 | 2611.40 ± 31.01 | 2808.39 ± 12.24 | 863.47 ± 42.42 | 3343.89 ± 49.49 | 2533.28 ± 35.45 | 2920.52 ± 15.11 | 2.13 ± 0.06 | 0.68 ± 0.01 | 1.32 ± 0.035 | 0.42 ± 0.004 |

#### Section S3: Adult shell height

Oysters sampled for histology were also measured for shell height using digital calipers (mm), defined as the maximum distance from the umbo along the dorsal/ventral axis. Shell height was compared between treatments prior to and after $pCO_2$ exposure using two-way Analysis of Variance (ANOVA) for each cohort. Shell height was also compared among cohort using one-way ANOVA, excluding the younger O-2 cohort due to their smaller initial size.

Prior to $pCO_2$ treatments, adult shell height did not vary between temperatures treatments, but did among F1 cohorts, with D smaller than F (p=0.043). After $pCO_2$ treatment, D was smaller than both F (p=3.4e-6) and O-1 (p=3.5e-6). The O-1 cohort increased in size (p=0.019) in ambient $pCO_2$, but not in high $pCO_2$. No size differences among pre-$pCO_2$ and post-$pCO_2$ treatments were observed in the F, D or O-2 cohorts.

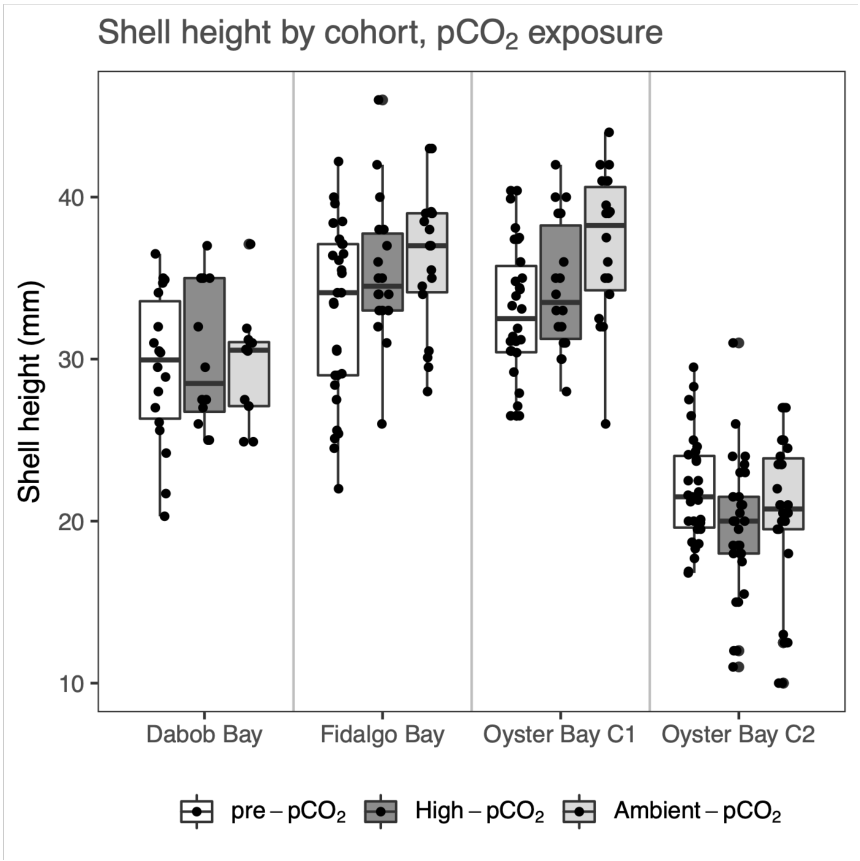


**Figure S1:** Adult shell height for each cohort, after temperature treatment but before $pCO_2$ treatment ("Pre-$pCO_2$"), and after 52 days in High-$pCO_2$ (3045±488 µatm, n=39), and Ambient-$pCO_2$ (7.82±0.02, n=39).

| **Table S2:** Gonad stage designations, adapted from da Silva 2009 | | |
|---|---|---|
| Stage | Designation | Description |
| 0 | Inactive | Empty follicles, no presence of male or female gonad tissue. |
| 1 | Early gametogenesis | Gametes were mostly attached to the follicle wall. In the male developing line, spermatogonia and spermatocytes, very few spermatids; in the female developing line, mostly attached, developing ovogonia and early ovocytes. |
| 2 | Advanced gametogenesis | In the male developing line few spermatogonia, spermatocytes and spermatid balls were dominant, and few spermatozoa balls appeared in the follicle lumen; in the female developing line, few ovogonia present, ovocytes in vitellogenesis but attached were dominant, and ovocytes in post-vitellogenesis and located free in the follicle lumen less abundant. |
| 3 | Ripe | In both male and female developing lines, follicles contained mature gametes and sometimes a thin layer of primary germ cells. Abundant spermatozoa balls and mature ovocytes filled the follicle lumen, in male line and female line, respectively. |
| 4 | Spawned (full or partial), and/or resorbing | Gametes had been released, and follicles are dilated but lumen was empty, or contained residual mature gametes; residual oocytes of various sizes were sparsely distributed; residual spermatids were dissociated within follicle lumen. In some cases phagocytes were observed within follicles to re-absorb residual gametes. In many cases residual gametes of one sex remained, while developing gametes of the other sex were abundant. |

| **Table S3:** Gonad sex designations, from da Silva 2009 | | | |
|---|---|---|---|
| Sex (uncollapsed, from da Silva) | Sex, collapsed for statistical analyses | Designation | Description |
| F | F | Female | Follicles contain only female gonad material (any stage) |
| HPF | F | Hermaphroditic, predominantly female | Follicles contain predominantly female but also some male gonad material |
| H | H | Hermaphroditic | Follicles contain approximately half male and half female gonad material |
| HPM | M | Hermaphroditic, predominantly male | Follicles contain predominantly male but also some female gonad material |
| M | M | Male | Follicles contain only male gonad material (any stage) |
| I | I | Indeterminate | Follicles are empty, collapsed, or only undifferentiated gonia are visible |

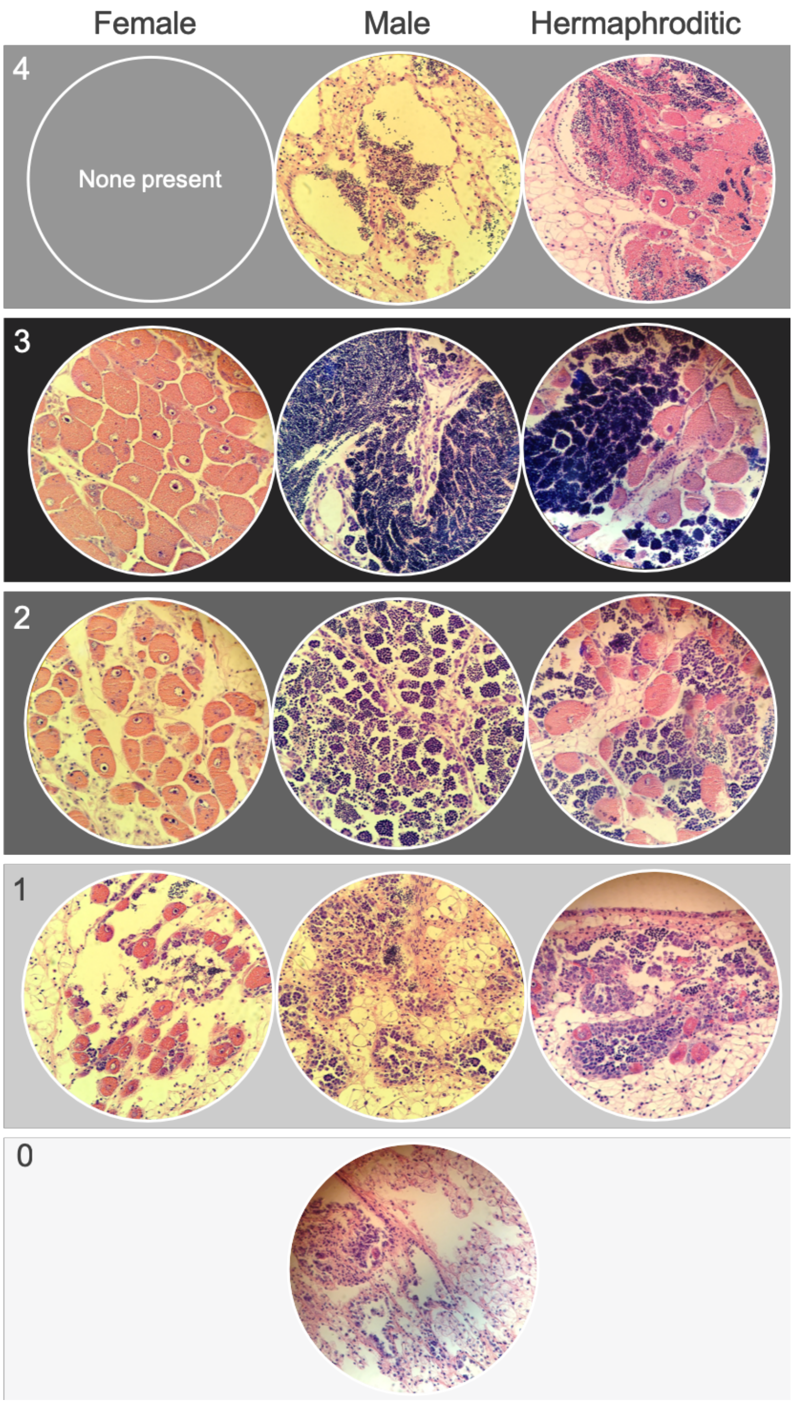


**Figure S2:** Examples of *Ostrea lurida* gonad stage designations. Stage 0 (no activity/sex differentiation); Stage 1 (early gametogenesis); Stage 2 (advanced gametogenesis); Stage 3 (late gametogenesis / ripe); Stage 4 (spawned and/or resorbing).

#### Section S4: Gonad sex and stage details and cohort traits

Of all sampled oysters, 53.4% were hermaphrodites, 8.3% contained only female gametes, and 31.1% contained only male gametes, and the remaining 7.2% were indeterminate. Across all treatments, gonad sex (collapsed for comparison) differed significantly among cohorts ($\chi^2$=55.8, p=1.0e-4). Fifty percent of all O-1 oysters sampled were female or hermaphroditic-primarily female (HPF), while 33%, 24% and 11% of D, F, and O-2 were female or HPF. Male or hermaphroditic-primarily male oysters comprised 29%, 48%, 59% and 69% of O-1, D, F, and O-2 cohorts, respectively.

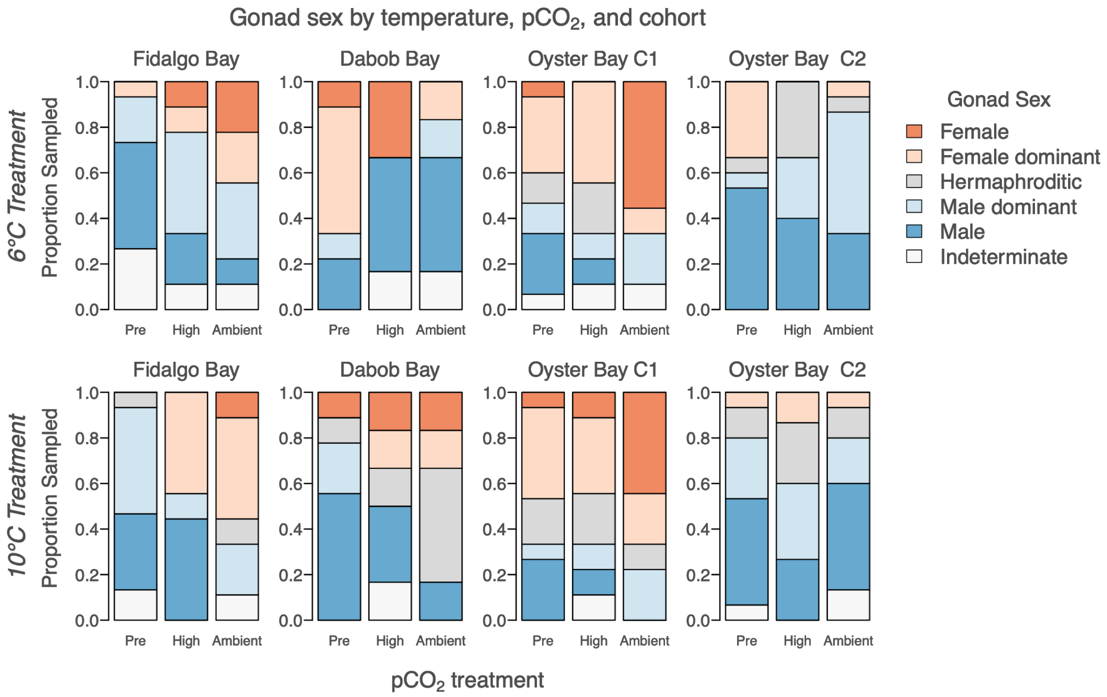


**Figure S3**: Gonad sex for each cohort, after temperature treatment but before $pCO_2$ treatment ("Pre"), and after 52 days in high $pCO_2$ (3045±488 µatm, n=39, "High"), and ambient $pCO_2$ (7.82±0.02, n=39, "Ambient"), separated by temperature treatment (6°C and 10°C).

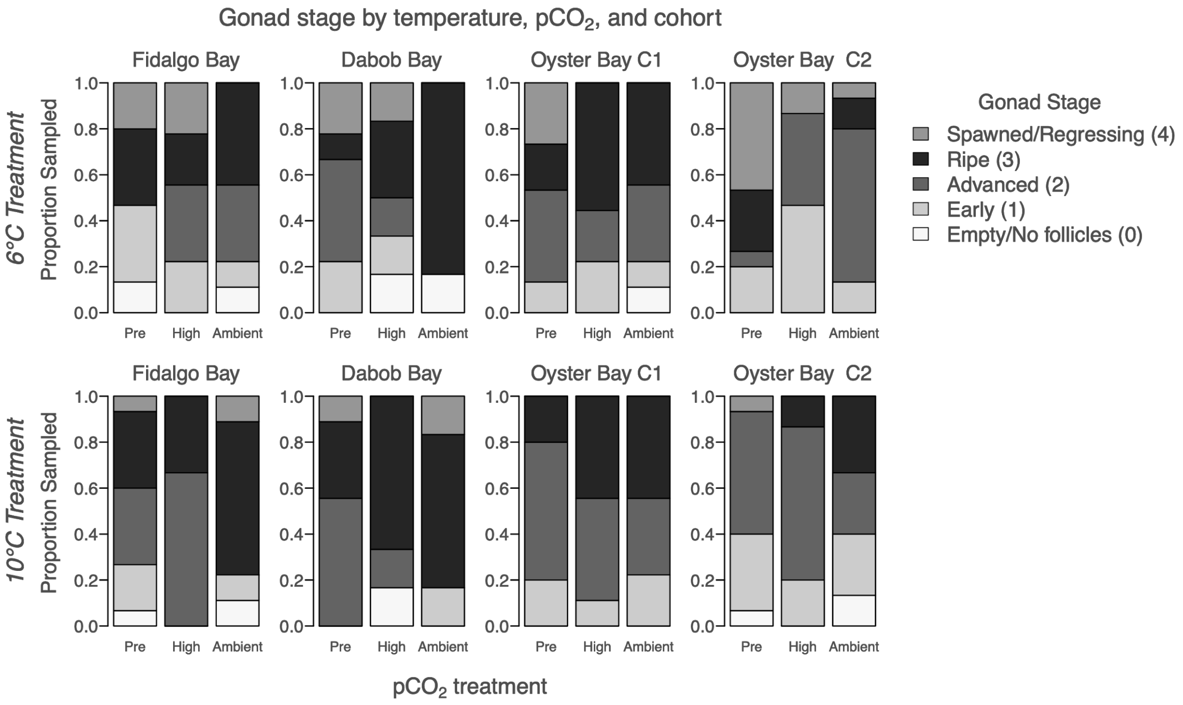


**Figure S4:** Gonad stage of the dominant sex for each cohort, after temperature treatment but before $pCO_2$ treatment ("Pre"), and after 52 days in high $pCO_2$ (3045±488 µatm, n=39, "High"), and ambient $pCO_2$ (7.82±0.02, n=39, "Ambient"), separated by temperature treatment (6°C and 10°C).

**Table S4**: Frequency of dominant gonad stages by cohort and $pCO_2$ exposure, separated by winter temperature treatment (6°C, 10°C). Results of Chi-Square between $pCO_2$ treatments are shown as p-values for dominant stage (Dom-stage), dominant sex (Dom-sex), Male stages (Male), and female stages (female). Across all cohorts in both 6°C and 10°C treatments, dominant stage and male stage differed by $pCO_2$ treatments. Stage differences were detected between pre-treatment (pre-$pCO_2$) and after treatment for ambient and high $pCO_2$ separately, and are indicated by superscripts A and B: A= pre-treatment and ambient $pCO_2$ differed; B=pre-treatment and high $pCO_2$ differed ($p<0.05$). Stage and sex comparisons in bold indicate that they were significantly different between temperature treatments, prior to $pCO_2$ treatments ("Pre" columns).

*Dominant gonad stage for 6°C treatment, by cohort and $pCO_2$ exposure*

| | *Fidalgo Bay* | | | *Dabob Bay* | | | *Oyster Bay C1* | | | *Oyster Bay C2* | | | *All cohorts*<br>***Dom-stage$^{A}$:*** *p=0.036*<br>*Dom-sex$^{A}$: p=0.202*<br>***Male$^{A,B}$:*** *p=0.030*<br>*Female: p=0.219* | | |
|---|---|---|---|---|---|---|---|---|---|---|---|---|---|---|---|
| *Gonad Stage* | Pre $pCO_2$ | High $pCO_2$ | Amb $pCO_2$ | Pre $pCO_2$ | High $pCO_2$ | Amb $pCO_2$ | Pre $pCO_2$ | High $pCO_2$ | Amb $pCO_2$ | Pre $pCO_2$ | High $pCO_2$ | Amb $pCO_2$ | Pre $pCO_2$ | High $pCO_2$ | Amb $pCO_2$ |
| Spawned / Resorbing (4) | 3 | 2 | 0 | 2 | 1 | 0 | 4 | 0 | 0 | 7 | 2 | 1 | *16* | *5* | *1* |
| Ripe (3) | 5 | 2 | 4 | 1 | 2 | 5 | 3 | 5 | 4 | 4 | 0 | 2 | *13* | *9* | *15* |
| Advanced (2) | 0 | 3 | 3 | 4 | 1 | 0 | 6 | 2 | 3 | 1 | 6 | 10 | *11* | *12* | *16* |
| Early (1) | 5 | 2 | 1 | 2 | 1 | 0 | 2 | 2 | 1 | 3 | 7 | 2 | *12* | *12* | *4* |
| Empty Follicles (0) | 2 | 0 | 1 | 0 | 1 | 1 | 0 | 0 | 1 | 0 | 0 | 0 | *2* | *1* | *3* |
| *Total Sampled* | *15* | *9* | *9* | *9* | *6* | *6* | *15* | *9* | *9* | *15* | *15* | *15* | *54* | *39* | *39* |

*Dominant gonad stage for 10°C treatment, by cohort and $pCO_2$ exposure*

| | *Fidalgo Bay* | | | *Dabob Bay* | | | *Oyster Bay C1* | | | *Oyster Bay C2* | | | *All cohorts*<br>***Dom-stage$^{A}$:*** *p=0.008*<br>*Dom-sex$^{A}$: p=0.73*<br>***Male$^{A}$:*** *p=0.020*<br>*Female: p=1* | | |
|---|---|---|---|---|---|---|---|---|---|---|---|---|---|---|---|
| *Gonad Stage* | Pre $pCO_2$ | High $pCO_2$ | Amb $pCO_2$ | Pre $pCO_2$ | High $pCO_2$ | Amb $pCO_2$ | Pre $pCO_2$ | High $pCO_2$ | Amb $pCO_2$ | Pre $pCO_2$ | High $pCO_2$ | Amb $pCO_2$ | Pre $pCO_2$ | High $pCO_2$ | Amb $pCO_2$ |
| Spawned / Resorbing (4) | 1 | 0 | 1 | 1 | 0 | 1 | 0 | 0 | 0 | 1 | 0 | 0 | *3* | *0* | *2* |
| Ripe (3) | 5 | 3 | 6 | 3 | 4 | 4 | 3 | 4 | 4 | 0 | 2 | 5 | *11* | *13* | *19* |
| Advanced (2) | 5 | 6 | 0 | 5 | 1 | 0 | 9 | 4 | 3 | 8 | 10 | 4 | *27* | *21* | *7* |
| Early (1) | 3 | 0 | 1 | 0 | 0 | 1 | 3 | 1 | 2 | 5 | 3 | 4 | *11* | *4* | *8* |
| Empty Follicles (0) | 1 | 0 | 1 | 0 | 1 | 0 | 0 | 0 | 0 | 1 | 0 | 2 | *2* | *1* | *3* |
| *Total Sampled* | *15* | *9* | *9* | *9* | *6* | *6* | *15* | *9* | *9* | *15* | *15* | *15* | *54* | *39* | *39* |

#### Section S5: Larval collection differences among cohorts

Total larvae collected differed by cohort ($F(3,8)=15.3$, $p=0.001$). O-1 produced significantly more total larvae than F and O-2 ($p=0.0094$, $p=0.0014$, respectively), and D produced more total larvae compared to O-2 ($p=0.022$). Total larvae released by O-1, F, D, and O-2 was 10.1M, 3.6M, 2.7M and 2.1M, respectively. The same patterns were observed in average daily larvae released by cohort ($F(3,20)=8.9$, $p=0.0009$). Date of first larval release differed by cohort ($F(3,8)=15.1$, $p=0.0012$). Oyster Bay cohorts (O-1 and O-2) released larvae 9.9 days earlier than F and D cohorts on average. Larval pulse frequency differed by cohort ($F(3,8)=9.8$, $p=0.0046$). On average, O-1, O-2, F, and D released larvae 6.4±2.3, 8.0±2.9, 3.8±1.9, and 2.8±1.0 days, respectively. The O-1 cohort released larvae more frequently than F ($p=0.017$), and O-2 more frequently than both F and D ($p=0.0066$, $p=0.043$, respectively).

**Table S5**: Timing and magnitude of larval production in 4 *Ostrea lurida* cohorts previously exposed to different winter temperatures (6°C and 10°C), then $pCO_2$ treatments ("Amb." is Ambient=841±85 µatm, pH 7.82±0.02, High=3045±488 µatm). Fidalgo Bay, Dabob Bay, and Oyster Bay are previously identified as genetically distinct populations (Jake E. Heare et al., 2017; J. Emerson Heare et al., 2018a). Two Oyster Bay cohorts were used (O-1, O-2), with O-2 being the offspring of O-1 and likely all siblings. For each metric, total ("Tot.") or mean ("Ave.") of all cohorts combined for each treatment is shown.

| Cohort | | *Fidalgo Bay (2 reps)* | | *Dabob Bay* | | *Oyster Bay - F1 (2 reps)* | | *Oyster Bay - F2* | | *All cohorts combined* | |
|---|---|---|---|---|---|---|---|---|---|---|---|
| $pCO_2$ treatment | | Amb. $pCO_2$ | High $pCO_2$ | Amb. $pCO_2$ | High $pCO_2$ | Amb. $pCO_2$ | High $pCO_2$ | Amb. $pCO_2$ | High $pCO_2$ | Amb. $pCO_2$ | High $pCO_2$ |
| No. of broodstock | 6°C | 15/14 | 14/15 | 14 | 15 | 15/16 | 17/17 | 117 | 126 | Tot: 191 | Tot: 204 |
| | 10°C | 15/14 | 14/14 | 9 | 16 | 17/17 | 15/15 | 115 | 111 | Tot: 177 | Tot: 185 |
| Date of first release | 6°C | 142/139 | 156/145 | 163 | 133 | 134/137 | 140/141 | 139 | 137 | Ave: 142 | Ave: 142 |
| | 10°C | 145/137 | 144/141 | 144 | 146 | 135/134 | 134/137 | 131 | 131 | Ave: 138 | Ave: 139 |
| Date of max release | 6°C | 149/146 | 161/145 | 163 | 168 | 135/156 | 141/165 | 154 | 143 | Ave: 151 | Ave: 154 |
| | 10°C | 145/137 | 144/144 | 151 | 147 | 135/152 | 151/137 | 140 | 143 | Ave: 143 | Ave: 144 |
| Date of last release | 6°C | 173/149 | 173/191 | 187 | 173 | 170/170 | 168/168 | 173 | 161 | Ave: 170 | Ave: 172 |
| | 10°C | 158/168 | 184/191 | 182 | 175 | 191/187 | 175/173 | 166 | 170 | Ave: 175 | Ave: 178 |
| Ave. daily larvae released ($x10^3$) | 6°C | 21/38 | 99/127 | 53 | 107 | 139/114 | 103/175 | 58 | 43 | Ave: 79 | Ave: 110 |
| | 10°C | 139/84 | 58/56 | 117 | 101 | 192/107 | 133/111 | 36 | 57 | Ave: 108 | Ave: 88 |
| Total larvae released ($x10^3$) | 6°C | 127/150 | 591/892 | 105 | 959 | 697/ 1,482 | 719/ 1,397 | 518 | 389 | Tot: 3.08M | Tot: 4.95M |
| | 10°C | 695/421 | 345/393 | 939 | 705 | 1,918/ 1,502 | 933/ 1,441 | 466 | 689 | Tot: 5.9M | Tot: 4.5M |
| Total larvae released per broodstock ($x10^3$) | 6°C | 2.4/3.0 | 11.7/16.5 | 2.5 | 21.3 | 12.9/25.7 | 11.7/22.8 | 2.0 | 1.4 | Ave: 6.6 | Ave: 13.5 |
| | 10°C | 1.3/8.4 | 6.8/7.8 | 34.8 | 14.7 | 31.3/24.6 | 17.3/26.7 | 1.8 | 2.8 | Ave: 18.8 | Ave: 11.7 |
| Maximum release ($x10^3$) | 6°C | 111/140 | 247/308 | 105 | 356 | 462/484 | 250/809 | 133 | 131 | Ave: 239 | Ave: 350 |
| | 10°C | 248/246 | 298/186 | 437 | 268 | 555/407 | 378/379 | 108 | 241 | Ave: 333 | Ave: 292 |
| No. big release days (>10k) | 6°C | 2/1 | 4/4 | 1 | 4 | 3/8 | 5/5 | 6 | 5 | Tot: 21 | Tot: 27 |
| | 10°C | 3/3 | 2/3 | 5 | 5 | 7/7 | 5/10 | 11 | 10 | Tot: 36 | Tot: 35 |

#### Section S6: Larval rearing methods and survival

##### Larval rearing methods

Larvae collected between May 19 and July 6 were separated by treatment and cohort and reared over 67 days from May 19 to July 25. For all culture tanks, seawater was heated to 18°C in a common 1,610-L recirculating reservoir (1610 L) using Aqua Logic digital temperature controllers (TR115SN), dosed with live algae cocktail via an Iwaki metering pump to achieve 100,000 cells/mL, and distributed to culture tanks. Larvae were grown in two connected 19-L flow-through tanks (19-L; 8-L/hr) with aerated, filtered seawater (1 µm) at 18°C. The two-tank larval system was used to cull dead larvae: water flowed from one 19-L tank where larvae were added but non-swimming larvae would remain ("mortality tank") to the next ("live tank"), carrying live, swimming larvae which were then contained on a 100 µm screen. Twice weekly, live larval tanks were screened into three size classes: 100 µm < X < 180 µm ("100 µm"), 180 µm < X < 224 µm ("180 µm"), >224 µm ("224 µm", which is when *O. lurida* larvae are near metamorphosis). Each size class was subsampled and counted, then the 100 µm and 180 µm classes returned to larval tanks. The number of live larvae returned to culture tanks informed stocking of newly released larvae. To maximize genetic diversity of offspring, newly spawned larvae (≲ 50,000) were added to culture tanks continuously to a maximum stocking density of 200,000 larvae (~10 larva/mL) (PSRF pers. communication & FAO manual). The contents in the mortality buckets were collected during biweekly screenings on a 100 µm screen to count live and dead oysters, but live were not kept.

During the twice weekly screening days, larvae that were larger than 224 µm were moved to downwelling setting silos, separated by cohort, temperature and $pCO_2$ treatment. Setting tanks were 180 µm silos with 18°C filtered seawater (1 µm) dosed with live algae, which then flowed into each silo from 8-L/hr irrigation drippers. Pacific oyster shell fragments (224 - 450 µm) were sprinkled into each silo to cover the surface to provide a settlement substrate. Silos were cleaned with freshwater (18°C) several times per week. Live, metamorphosed oysters were counted on August 12 for survival rate from 224 um to post-metamorphosis ("post-set"), then transferred to 450 µm silos with ~17°C upwelling filtered seawater (5 µm) to continue rearing. Oysters were fed live algae using a gravity algae header tank, and rinsed 1-2 times per week with freshwater. On October 4, when oysters were between 13-20 weeks old, all groups were moved to screen pouches separated by cohort x temperature x $pCO_2$, affixed to the inside of shellfish cages, and hung in Clam Bay until June 2018.

**Table S6**: Tank densities during offspring rearing.

| | | *Fidalgo Bay (2 replicates)* | | *Dabob Bay* | | *Oyster Bay - F1 (2 replicates)* | | *Oyster Bay - F2* | | *All* | |
|---|---|---|---|---|---|---|---|---|---|---|---|
| | | Amb pCO2 | High $pCO_2$ | Amb pCO2 | High $pCO_2$ | Amb pCO2 | High $pCO_2$ | Amb pCO2 | High $pCO_2$ | Amb pCO2 | High $pCO_2$ |
| Cumulative larvae stocked | 6°C | 169,475 | 353,916 | 65,667 | 255,371 | 477,403 | 532,417 | 353,769 | 179,717 | 1,066,314 | 1,321,421 |
| | 10°C | 319,277 | 271,016 | 283,858 | 310,875 | 937,920 | 660,451 | 324,166 | 237,790 | 1,865,221 | 1,480,132 |
| Mean, larvae density (in 20L tank) | 6°C | 29,148 ± 33,256 | 54,265 ± 44,431 | 22,000 ± 22,036 | 43,293 ± 43,168 | 50,260 ± 51,338 | 58,662 ± 64,276 | 47,231 ± 38,553 | 31,877 ± 30,341 | 39,270 ± 40,579 | 46,777 ± 47,718 |
| | 10°C | 40,150 ± 47,430 | 29,486 ± 30,781 | 49,584 ± 33,070 | 56,081 ± 49,214 | 88,962 ± 71,752 | 74,562 ± 72,312 | 53,588 ± 50,765 | 32,423 ± 41,853 | 60,478 ± 5,7472 | 48,774 ± 54,447 |
| Median larvae density | 6°C | 18,013 | 42,783 | 19,117 | 26,160 | 33,797 | 32,608 | 45,667 | 20,800 | 26,397 | 28,192 |
| | 10°C | 16,000 | 20,650 | 50,317 | 45,833 | 83,577 | 48,800 | 29,833 | 7,877 | 50,317 | 34,935 |
| Total eyed larvae | 6°C | 11,119 | 11,780 | 2,496 | 10,686 | 11,931 | 6,029 | 22,186 | 9,735 | 47,732 | 38,230 |
| | 10°C | 3,737 | 2,978 | 13,862 | 19,815 | 59,929 (split) | 10,670 | 13,828 | 2,910 | 91,356 | 36,373 |
| Total post set (singles) | 6°C | 1,503 (split) | 670 | 501 | 834 | 124 | 192 | 356 | 334 | 2,484 | 2,030 |
| | 10°C | 626 | 75 | 1311 | 1091 | 52 | 34 | 246 | 113 | 2,235 | 1,313 |
| Juvenile shell height (mm) | 6°C | 9.0±2.7 | 8.4±3.5 | 6.5±1.9 | 4.9±2.3 | 11.2±3.5 | 11.0±3.4 | 11.0±3.7 | 7.5±3.0 | | |
| | 10°C | 9.7±3.0 | 12.3±5.4 | 7.0±2.3 | 6.6±2.9 | 11.0±3.8 | 12.2±4.2 | 10.7±3.7 | 13.4±4.4 | | |
| Juveniles deployed | 6°C | 240 | 257 | 240 | 240 | 85 | 77 | 122 | 90 | 677 | 664 |
| Juveniles survived | 6°C | 60 | 159 | 72 | 81 | 32 | 45 | 22 | 4 | 186 | 289 |

**Larval survival estimates**
Larval survival was estimated from both twice-weekly larval counts and cumulative survival counts. Percent survival between biweekly larval counts was calculated by summing the number of live larvae in all size classes (100 µm, 180 µm, 224 µm), dividing by the number of live larvae restocked after the previous count, plus all new larvae added since. Cumulative percent survival from newly released larvae ("new larvae") to the near-metamorphosis stage ("eyed larvae"), and to post-metamorphosis ("post-set") were compared between treatments based on total number of new larvae stocked in culture tanks and eyed larvae in setting tanks over the larval rearing period. During larval rearing, culture tank densities were capped at 200,000 larvae (~10 larvae/mL), but ranged during the 67 day larval rearing period due to varying mortality and larval release timing. Daily tank densities were estimated from twice-weekly larval counts and number of new larvae added, then compared between temperature x $pCO_2$ treatments using a Kruskal-Wallis Test.

Biweekly larval survival, cumulative survival from new to eyed larvae, and survival from eyed larvae to post-set were compared among cohort x temperature x $pCO_2$ treatments using ANCOVA on fitted linear regression models. For biweekly percent survival, square-root arcsine transformation was applied, and biweekly tank density was included as a random effect. For cumulative survival models, mean stocking density and total larvae stocked in culture tanks were examined as candidate random effects with Pearson's correlation using `pairs` and `cor`. For post-set survival, cumulative eyed larvae stocked in setting tanks and percent survival to eyed larvae stage were also tested and survival data was log-transformed. Tank density factors that correlated significantly with cumulative survival were considered as random effects in full regression models alongside cohort, temperature and $pCO_2$. All models were optimized using stepwise deletion and selected based on AIC value, adjusted R-squared, and F-statistic.

**Larval survival results**
Larval survival between biweekly counts did not differ by $pCO_2$ or temperature, but did differ by cohort (F(3,230)=5.73, p=8.5e-4). Pairwise tests indicate that O-1 survival was significantly lower than D (p=3.8e-4), O-2 (p=5.4e-4), and F (p=0.019). Mean biweekly survival of D, F, O-2, and O-1 cohorts was 62±22%, 59±24%, 55±24%, and 49±28%, respectively. Cumulative survival from new- to eyed-larvae was low across all treatments, and did not differ by parental temperature treatment (F(1,14)=2.3, p=0.15), parental $pCO_2$ (F(1,14)= 1.9, p=0.19), or cohort (F(3,12)= 1.4, p=0.29) (Table 3). Cumulative survival from eyed larvae to post-set) ranged from 0.2% to 26.5% and differed by cohort (F(3,11)=3.8, p=0.04). Pairwise tests revealed that this was influenced by low survival in the O-1 group and significance was not strong after removing O-1 (F(2,9)=4.1, p=0.06). No survival differences through metamorphosis were detected between $pCO_2$ or temperature treatments.

Tank density prior to each biweekly screening was a significant factor influencing survival between bi-weekly counts (F(1,230)=10.4, p=0.0015) and therefore was included as a random effect in the biweekly survival regression model. Mean stocking densities across the 67-day rearing period in O-1, D, O-2, and F were 76,500±71,100, 54,400±424000, 47,000±46,200, and 43,500±42,700, respectively. No random effects were retained in the cumulative survival from new- to eyed-larvae model. Total larvae stocked in larval culture tanks correlated with survival from eyed-larvae to post-set (i.e. through metamorphosis), and therefore was included as a random effect in the post-set survival model.

The number of days between first and last larval collection, and first and last eyed larvae varied by cohort, although this was not significant. Across treatments, eyed larvae were present soonest in F (14.5±2.5 days), followed by O-1 (16.5±1.75 days), O-2 (17.25±1.25 days), and lastly D (18.25±3 days) ($F(3,12)=2.0$, $p=0.16$). The number of days between stocking the last batch of newly released larvae, and collecting the last eyed larvae were 22±5.8, 23.25±7.4, 29.5±4.7, and 32±4.8 for O-1, F, D, and O-2, respectively.

**Table S7**: Larval survival estimates by parental treatment and cohort.

*Larval survival, by treatment and cohort*

| | | *Fidalgo Bay* | | *Dabob Bay* | | *Oyster Bay - F1* | | *Oyster Bay - F2* | | *All cohorts* | |
|---|---|---|---|---|---|---|---|---|---|---|---|
| | | Amb $pCO_2$ | High $pCO_2$ | Amb $pCO_2$ | High $pCO_2$ | Amb $pCO_2$ | High $pCO_2$ | Amb $pCO_2$ | High $pCO_2$ | Amb $pCO_2$ | High $pCO_2$ |
| Average biweekly larval survival | 6°C | 56±25% | 62±26% | 69±18% | 49±21% | 44±23% | 52±27% | 60±24% | 59±28% | 56±24% | 56±26% |
| | 10°C | 50±18% | 51±26% | 58±25% | 70±18% | 49±30% | 46±29% | 63±25% | 55±21% | 55±25% | 56±25% |
| Cumulative survival to eyed larvae | 6°C | 8.5% | 3.0% | 5.1% | 4.7% | 2.7% | 1.2% | 6.7% | 5.7% | 5.7±2% | 3.6±2% |
| | 10°C | 1.3% | 1.4% | 5.2% | 6.6% | 4.2% | 0.7% | 4.4% | 1.0% | 3.8±1% | 2.4±3% |
| Cumulative survival, eyed larvae to post set | 6°C | 13.8% | 5.9% | 26.5% | 9.3% | 1.1% | 3.6% | 1.7% | 3.5% | 10.8 ±10% | 5.6 ±2.7% |
| | 10°C | 18.5% | 2.7% | 9.7% | 6.0% | 0.2% | 0.7% | 1.9% | 5.8% | 7.6±8% | 3.8±3% |

#### Section S7: Juvenile survival and environmental data

Associations among juvenile oyster survival and environmental summary statistics were explored to evaluate factors that best explain spatial variation in oyster survival. The mean and standard deviation of each environmental variable (temperature, dissolved oxygen, salinity, chlorophyll, and pH) during deployment were assessed independently by binomial generalized linear mixed models (glmm) using glmer from the lme4 package (vs. 1.1-19), and Wald tests with type II error. Significant single-factor variables were included in a full model, then backwards deletion was used to identify significant environmental factors in the most parsimonious model. Significant variables predicting juvenile survival included mean temperature, mean pH, and dissolved oxygen standard deviation. Figure S6 show proportion survival by each environmental summary statistic. See the “07_Juvenile-deployment.R “ R code in the Spencer et al. 2019 GitHub repository for analysis details and results.

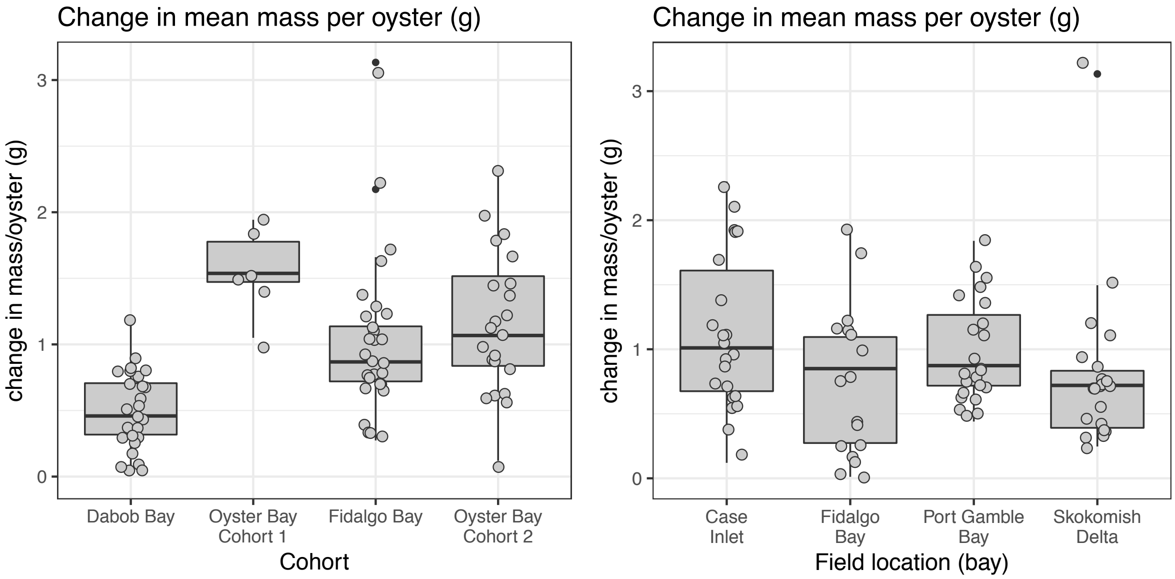


**Figure S5**: Juvenile mass change during field trial was significantly less in the Dabob Bay cohort than other 3 cohorts (left), and was significantly less in Fidalgo Bay than in Port Gamble Bay and Case Inlet (right). Mean mass / oyster represents the average final mass per oyster minus average initial mass within each deployment bag.

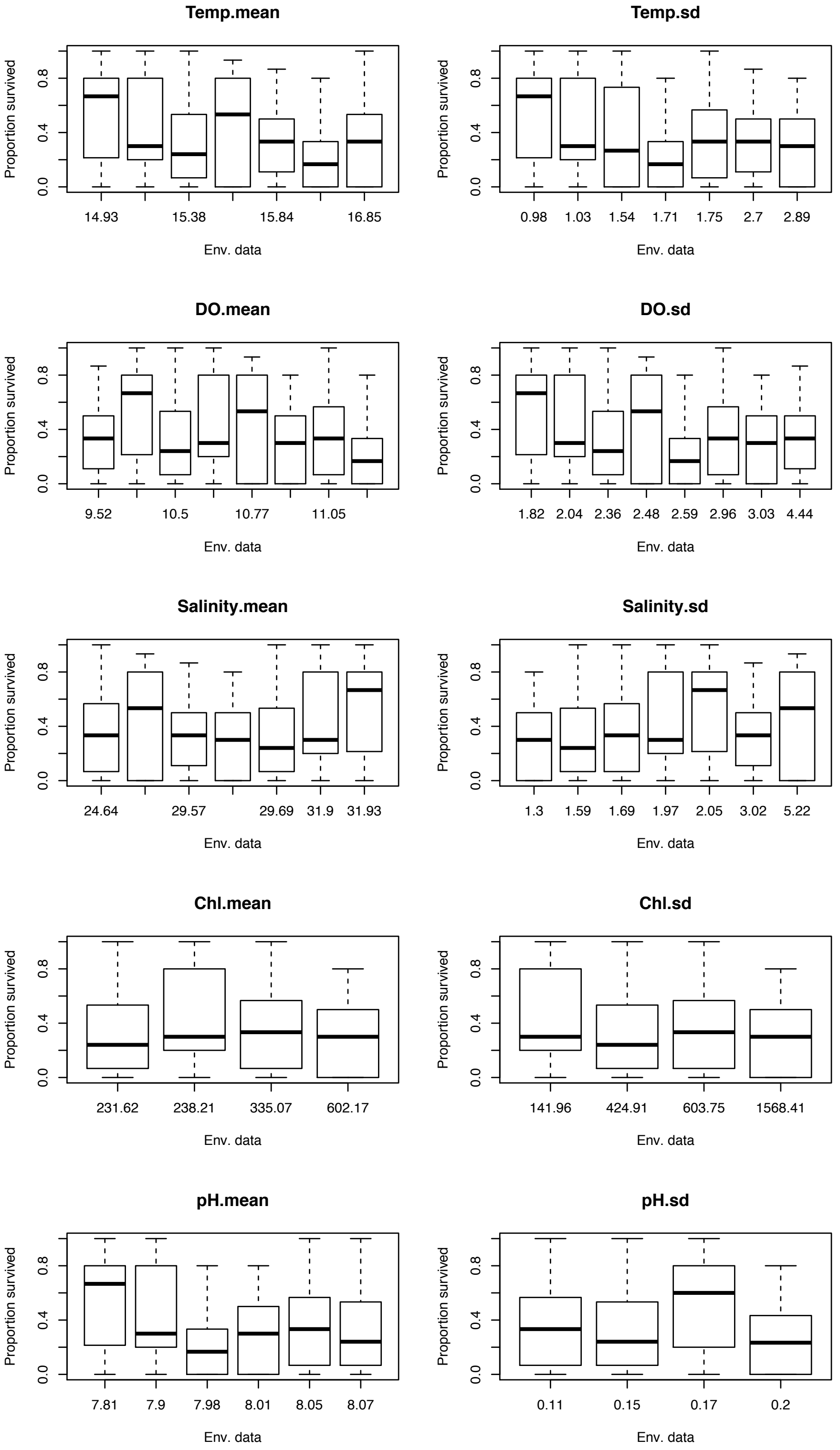


**Figure S6**: Juvenile proportion survival ~ environmental summary statistics. Model selection using backwards deletion indicates that survival was significantly related to mean temperature (“Temp.mean”), mean pH (“pH.mean”), and dissolved oxygen standard deviation (“DO.sd”).
